# Cysteines are critical determinants of spontaneous and seeded tau aggregation in cells

**DOI:** 10.64898/2026.03.12.711133

**Authors:** Parvathy Jayan, Simran Rastogi, Vaibhav Bommareddy, Chad M. Dashnaw, Jaime Vaquer-Alicea, Binh An Nguyen, Lorena Saelices, Sarah Shahmoradian, Charles L. White, Marc I. Diamond, Lukasz A. Joachimiak

**Affiliations:** Center for Alzheimer’s and Neurodegenerative Diseases, Peter O’Donnell Jr. Brain Institute, University of Texas Southwestern Medical Center, Dallas, TX 75390, USA; Graduate Program in Neuroscience, University of Texas Southwestern Medical Center, Dallas, TX, 75390, USA; Department of Biophysics, University of Texas Southwestern Medical Center, Dallas, TX 75390, USA; Department of Pathology, University of Texas Southwestern Medical Center, Dallas, TX, 75390, USA; Department of Biochemistry, University of Texas Southwestern Medical Center, Dallas, TX 75390, USA

## Abstract

The frontotemporal dementia-linked S320F mutation in the microtubule-associated protein tau promotes spontaneous aggregation, yet the structural basis of its amyloidogenesis remains unclear. Using cryo-electron microscopy, we determined the structure of an S320F_295-330_ tau fibril composed of parallel chains stabilized by the ^306^VQIVYK^311^ amyloid motif, with S320F buried in the fibril core and a C322-C322 disulfide linking two protofilaments. Although cysteines are dispensable for fibril formation by isolated peptide fragments in vitro, tau repeat domain constructs containing both C291 and C322 generate more potent seeds in cellular assays. In contrast, the C322S mutation suppresses spontaneous aggregation of S320F tau in cells, and combined C291S and C322S mutations inhibit seeded aggregation in both wild-type and S320F contexts. Systematic alanine mutagenesis coupled with seeding by tauopathy-derived material identifies cysteine residues as critical determinants of tau seeding, comparable in importance to core amyloid motifs. Together, these findings establish cysteines as central chemical regulators of tau aggregation and propagation.

## Introduction

Tau is an intrinsically disordered protein that was first discovered as a factor that regulates microtubule assembly^1^. Misfolding of tau into β-sheet rich amyloids is linked to several diseases, including Alzheimer’s disease (AD), frontotemporal dementia (FTD), corticobasal degeneration (CBD), progressive supranuclear palsy (PSP), commonly known as tauopathies^2^. Tau consists of an N-terminal domain, a proline-rich region, microtubule binding repeats (R1, R2, R3, and R4), and a C-terminus^2^. There are six isoforms of tau present in the central nervous system^3^, which differ in the presence or absence of N-terminal inserts and microtubule binding domain R2^4^. Cryo-EM studies on tau filaments obtained from the brains of patients with different tauopathies have revealed that distinct folds are associated with different diseases, which signifies a direct relationship between fibril conformation and disease^5–8^. Induction of tau aggregation *in vitro* is accelerated by coincubation with polyanions such as heparin or RNA^9–11^, and the resultant fibrils yield structures different from those observed in patients with tauopathies^12,13^. Recent data also suggests that phosphomimetic mutations can promote tau aggregation at high concentrations that do not require additional inducers^14^. The mechanisms by which a protein spontaneously adopts a defined amyloid conformation linked to disease remains a big question in the field.

The prion hypothesis posits that protein structure encodes disease phenotype and has been especially powerful in explaining the molecular diversity of human tauopathies^15–18^. Efforts are underway to leverage this structural complexity for more precise diagnostics by developing approaches that classify patients based on the conformations of pathological tau assemblies. As the catalog of tau amyloid structures expands, general principles of amyloid stabilization are emerging, with core motifs acting as dominant drivers of stability through burial of extensive nonpolar surfaces^19–25^. Systematic manipulation of sequence and assembly conditions is beginning to reveal how tau adopts disease-associated folds and, in some cases, forms assemblies spontaneously^14,26–29^. These observations raise fundamental questions: must a pathogenic tau conformation propagate in a prion-like manner to cause disease, or can cell-autonomous aggregation alone be sufficient? Moreover, are the rules governing spontaneous self-assembly distinct from those that control seeded aggregation in cells and organisms? Most tauopathies cases, including in AD, are sporadic, whereas mutations in the *MAPT* gene encoding microtubule-associated protein tau are linked to FTD, a group of neurodegenerative disorders marked by progressive degeneration of the frontal and temporal cortex^30–33^. To date, more than 50 disease-associated *MAPT* mutations have been identified, primarily within exons 9-12 that encode the tau repeat domain^34^. FTD-linked mutations, including P301L, P301S, R406W, V337M, and ΔK280, enhance tau aggregation propensity but still require exogenous inducers to initiate amyloid assembly^28,35–39^. Cryo-EM structures of P301S tau filaments from transgenic models, and more recently from FTD patients carrying P301T/L mutations, suggest that the wild-type proline at position 301 restricts conformational flexibility, thereby exerting a protective effect^8^. Similarly, cryo-EM studies of tau filaments extracted from brains of individuals harboring V337M and R406W mutations show paired helical and straight filament folds that closely resemble those seen in AD, consistent with ex vivo assemblies of tau fragments (297-391) bearing the V337M mutation^39^. Together, these findings highlight the importance of understanding how FTD-linked mutations modulate tau assembly to enable new therapeutic strategies.

In contrast to other *MAPT* mutations, the S320F substitution is the only known FTD-associated mutation^40^ to date that reported spontaneous aggregation of full-length tau and tau repeat domain, both in vitro and in cells^28,41^. Computational modeling revealed that mutation of serine to phenylalanine at position 320 reshapes the tau conformational ensemble by altering nonpolar residue clustering, thereby increasing exposure of amyloid-forming sequences^28^. Combining P301L with S320F in full-length tau leads to self-assembly in neuronal culture^42^ and animal models^43^. Fibril structures of tau amyloids isolated from FTD-tau patients harboring the S320F mutation reveals a fibril fold similar to the conformation observed in Picks disease (PiD)^44^. Indeed, modeling of the S320F mutation or similarly sized nonpolar residues in CBD and PiD improved predicted stability of the fibril and enhanced aggregation propensity^28^. These results demonstrate that transient interactions between nonpolar residues can shield amyloid motifs and suppress aggregation and that disruption of these interactions by a disease-associated mutation can stabilize alternative non-polar interactions that can drive self-assembly. Consistent with this model, mutation of nonpolar residues that interact with S320F to polar amino acids suppresses spontaneous aggregation in vitro and in cells^28^ and supported by the new FTD-tau S320F fibril structure^44^. Thus, subtle rewiring of transient interactions within intrinsically disordered proteins can have profound consequences for aggregation behavior. Although we proposed a mechanistic model for how S320F promotes tau assembly, the FTD-tau patient-derived amyloid structure adopted by S320F does not explain how this mutant uniquely enables spontaneous tau amyloid formation.

Here, we used cryo-EM to determine the structure of a self-aggregating S320F tau fragment spanning residues 295-330 (herein S320F_295-330_) to elucidate how this mutation drives spontaneous aggregation. We identify unusual structural features in which two monomers assemble in parallel, in-register alignment stabilized by an intermolecular disulfide bond. Surprisingly, mutation of the cysteine within S320F_295-330_ further enhances aggregation and increases seeding efficiency in cellular tau aggregation models. By contrast, in vitro aggregation of the tau repeat domain (tauRD) encoding S320F reveals that constructs retaining both cysteines (C291 and C322) exhibit more robust cellular seeding than single or double cysteine-to-serine mutants. In cellular contexts, mutation of cysteine 322 to serine in the context of tauRD S320F suppresses the spontaneous aggregation phenotype observed in cells. Moreover, wild-type tauRD or tauRD S320F harboring the double cysteine-to-serine mutations inhibits seeded aggregation across diverse seed sources, ranging from in vitro generated fibrils to tauopathy patient-derived material. To systematically assess the contribution of cysteine residues relative to other positions in tau, we performed multiplexed alanine scanning across the sequence and quantified seeding responses against AD, CBD and PSP brain extracts. We find that alanine substitutions at C291 and C322 exert pronounced, context-dependent effects on seeded aggregation comparable in magnitude to mutations within core amyloidogenic motifs. Together, these data support a central role for cysteine residues in regulating both spontaneous and seeded tau aggregation in vitro and in cells.

## Results

### Cryo-EM fibril structure of the S320F_295-330_ tau fragment

Prior work from our lab established a minimal fragment of tau spanning the R3 repeat and the C-terminal regulatory element of R2 in which the wild-type sequence did not aggregate yet the sequence encoding the S320F aggregated rapidly^28^. Using this tau fragment spanning residues 295-330 and encoding the S320F mutation (Fig. 1a, herein S320F_295-330_), we sought to determine the amyloid structure to understand how this mutation may promote tau assembly. S320F_295-330_ filaments were formed by incubating 300 µM peptide in 1x phosphate-buffered saline (PBS) (pH 7.4) and 2mM TCEP at 37°C. Fibril formation was confirmed by Thioflavin T (ThT) fluorescence and negative-stain TEM (Fig. 1b). Cryo-EM grids were optimized to produce dispersed fibrils suitable for cryo-EM data analysis. 6,509 movies were collected, and 102,937 filaments were picked manually. Particles were then extracted and classified using Relion^45^ to identify a dominant 2D class (Supplementary Fig. 1). These filaments exhibited a cross-over distance of 330 Å, corresponding to a twist angle and helical rise of -2.38° and 4.76 Å, respectively (Fig. 1c). Refinement of this model led to a final map with 3.7 Å resolution with well-defined features consistent with aromatic residues enabling unambiguous assignment of the sequence into the density using COOT^46^ and the atomic structure was further refined using Phenix^47^. The final model of the filament reveals a core comprised of three chains, all adopting unique conformations of the sequence (chain A, B, and C). Chains A and B span the entire sequence of R3 from residues 305/306 to 330, while chain C contains residues from 304 to 314 (Fig. 1d,e). Chains A and B interact with each other in parallel through a network of hydrophobic interactions at the N-terminus mediated by ^306^VQIVYK^311^. Chain C consists of residues from 304 to 314 that include the ^306^VQIVYK^311^ amyloid motif and interacts with the C-terminal end of chain A. The S320F mutation is also present in the fibril core, which indicates that not only the amyloid motif ^306^VQIVYK^311^, but also the S320F mutation are part of the amyloid core. By virtue of the parallel orientation of chains A and B, the ^306^VQIVYK^311^ and S320F interact with their corresponding matching sequence. From the cryo-EM structure of S320F_295-330_, we also found that there is an intermolecular disulfide bond between C322 of chain A and chain B. A study from Mandelkow et al from 1995^48^, first suggested that tau assembly may proceed through a disulfide dimer intermediate but no tau amyloid structures to date have shown an intermolecular disulfide bond in the reconstruction. While disulfide bonds play an important role in globular protein folding and stability, particularly in oxidizing environments, it is not clear what their role is in intrinsically disordered proteins and how they influence protein assembly towards amyloid states.

**Figure 1.**
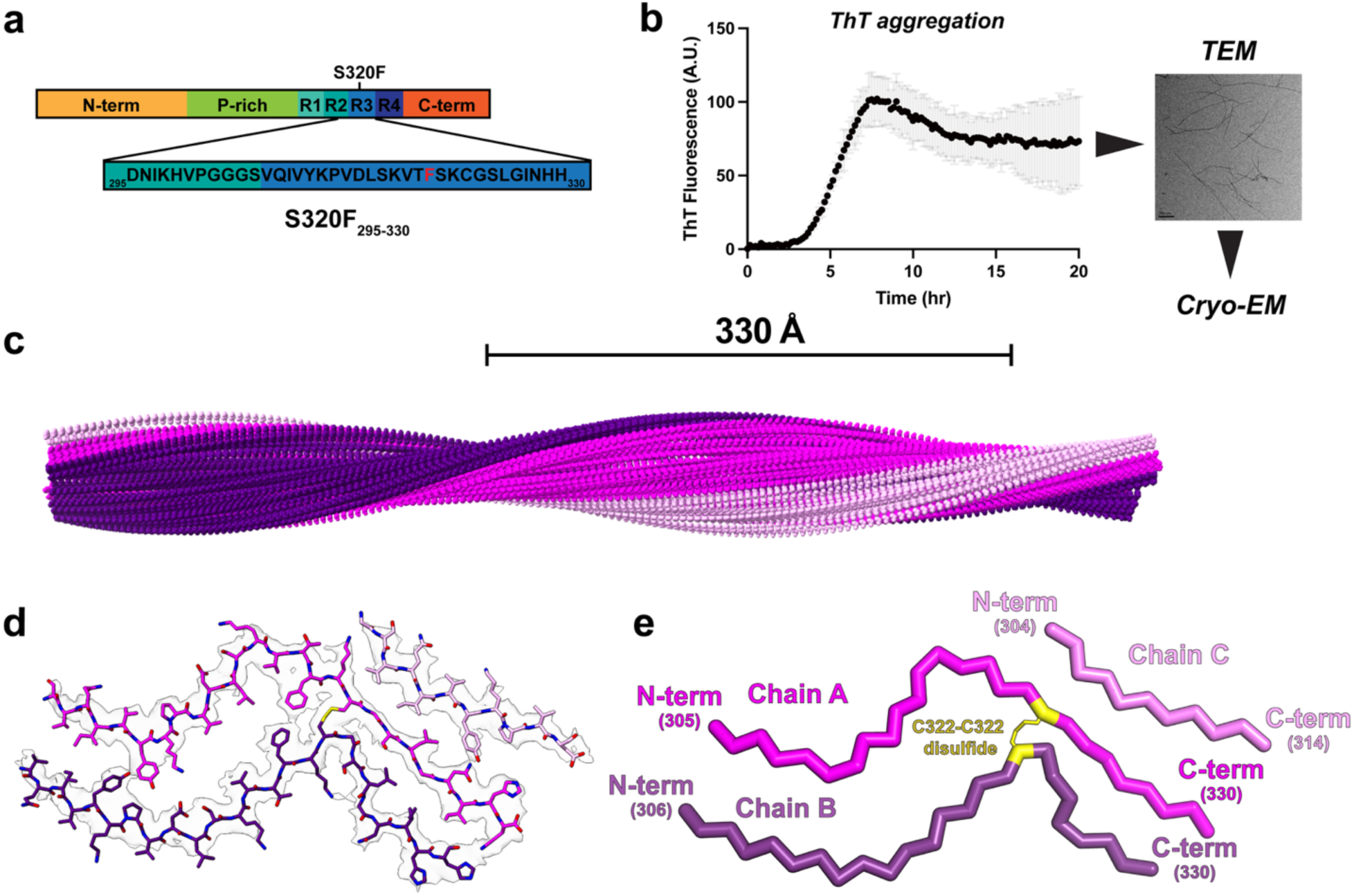
Structure of the tau S320F_295-330_ amyloid fibril. (**a**) Schematic representation of the S320F_295-330_ peptide fragment. Mutation of serine at 320 position is indicated (red) in the sequence. (**b**) ThT fluorescence aggregation assay of S320F_295-330_ in the presence of TCEP, reaction was carried out at 300 μM and is shown as an average across three technical replicates. Fibril formation was confirmed using TEM. (**c**) 3D reconstruction of the helical fibril illustrating the cross-over distance of 330 Å. The structure is shown as a spacefill model and colored by chain (pink, chain A; purple, chain B; light pink, chain C). (**d**) Cryo-EM density map and atomic model of S320F_295-330_ tau amyloid. Structure is shown as sticks with backbone and side chains. Map is contoured at 0.006. (**e**) Ribbon diagram of the S320F_295-330_ tau amyloid structure illustrating the parallel topology of chains A and B, and location of the C322-C322 interchain disulfide. The chains are shown as ribbons and colored separately (pink, chain A; purple, chain B; light pink, chain C). The disulfide is shown as sticks and is colored in yellow. The start and end residue positions for each chain are labeled.

### In silico thermodynamic profiling of S320F_295-330_ amyloid structure reveals interactions that stabilize the fibril

To quantify the energetic contributions of individual residues in our new structure to fibril stability, we performed in silico alanine-scanning mutagenesis using a validated framework for estimating the thermodynamic stability of amyloid fibrils^22^. This approach calculates residue-specific energetic changes (ΔΔG) between wild-type and alanine-substituted variants, providing a thermodynamic map of the fibril core. We compared ΔΔG values for each residue across the three fibril chains and as a function of 3-, 6-, 9-, and 12-layer assemblies (Fig. 2a,b). This analysis identified the ^306^VQIVYK^311^ amyloid motif at the N-terminus of chains A and B as a major determinant of fibril stability, with V306, I308, and Y310 contributing most strongly. Residues at the C-termini of chains A and B also contributed substantially to stability. Notably, mutation of I328 to alanine strongly destabilized the fibril, consistent with our previous observation that I328S inhibits aggregation of S320F_295-33028_. In contrast, mutation of F320 to alanine was predicted to be stabilizing, suggesting that the bulky phenylalanine side chain may be locally overpacked in chains A and B and that this strain is compensated by the intermolecular C322-C322 disulfide bond (Fig. 2c,d).

**Figure 2.**
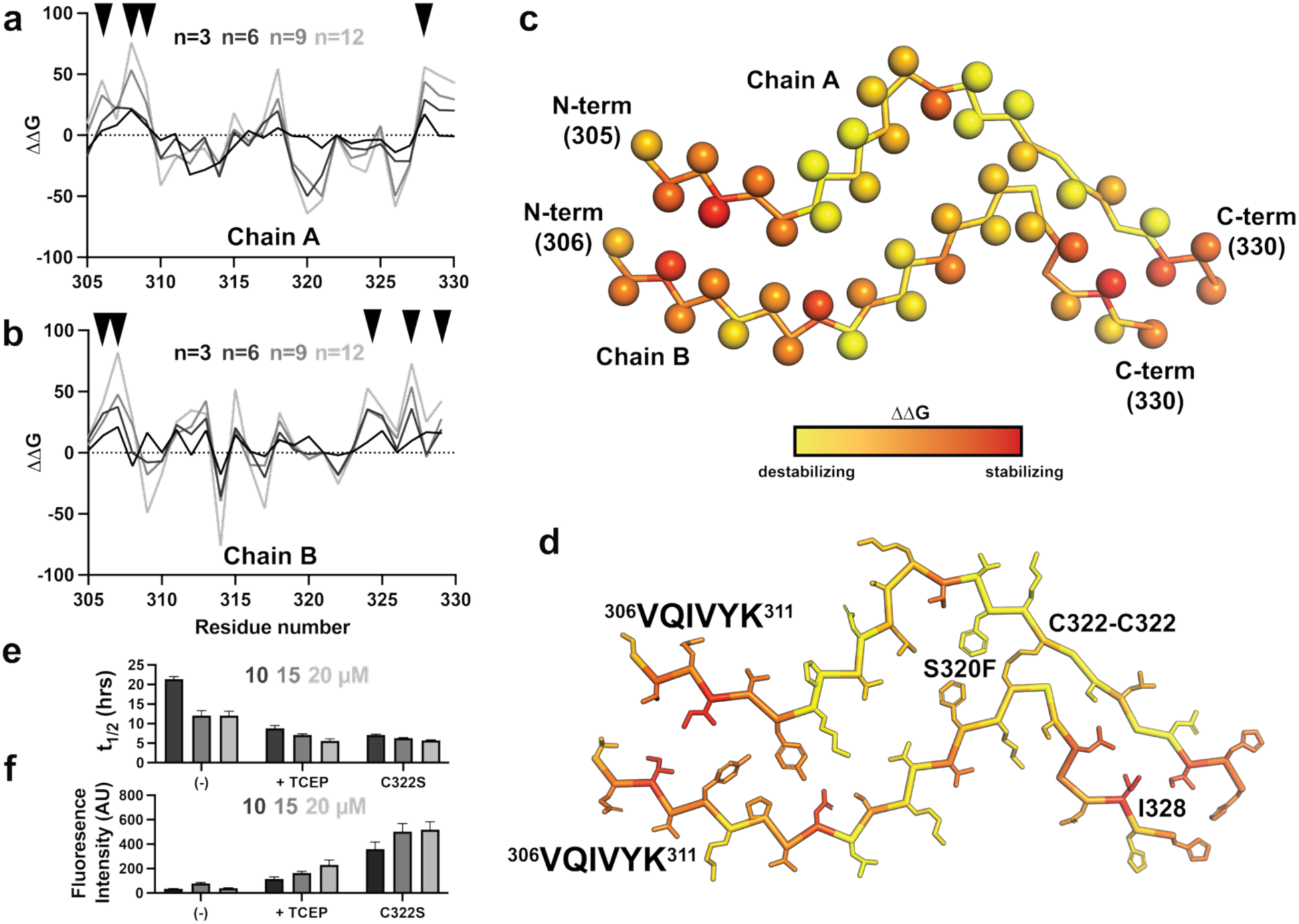
Thermodynamic profiling uncovers key features stabilizing S320F_295-330_ amyloid fibril. Calculated ΔΔG value of each amino acid in chain A (**a**) and chain B (**b**) calculated from in silico alanine mutagenesis for assemblies with increasing numbers of layers n=3, 6, 9 and 12. The ΔΔG values are colored from black (n=3) to grey (n=12). (**c**) Mapping per residue ΔΔG values onto chains A and B of the S320F_295-330_ cryo-EM structure uncovers hotspot residues in chains A and B at the N-terminus and C-terminus (>1 SD above the mean). Structures are shown as a single layer with the backbone shown as ribbons and the C-β of each residue is shown as a sphere. Each amino acid is colored by calculated ΔΔG from n=9 layer analysis. (**d**) The per residue ΔΔG values are shown mapped on a model of the S320F_295-330_ amyloid to highlight residues that stabilize the amyloid fold: (1) VQIVYK-VQIVYK, (2) C322-C322 disulfide and (3) I328-I328. Backbone is shown as ribbon and side chains are shown as sticks. (**e-f**) ThT fluorescence aggregation assay of S320F_295-330_ (-), S320F_295-330_ (TCEP), S320F_295-330_ C322S carried out at 10 μM (white), 15 μM (light grey) and 20 μM (grey). Data is shown as t_1/2max_ **(e)** and ThT fluorescence amplitude **(f)** parameters derived from a nonlinear regression fit to each aggregation curve. Data is presented as an average of three replicates with error bars as standard error.

To test the functional importance of this disulfide bond, we examined how redox conditions and cysteine substitutions affect fibril formation kinetics. Self-assembly of S320F_295-330_ was monitored under non-reducing conditions, in the presence of TCEP, or with C322 mutated to serine. Aggregation was followed for 24 h at 10, 15, and 20 µM using ThT fluorescence aggregation assays, and fibril formation was confirmed by negative-stain TEM at the endpoint (Supplementary Figs. 2a-c and 3). All samples exhibited concentration-dependent aggregation, with faster kinetics at higher peptide concentrations. In non-reducing conditions, S320F_295-330_ aggregated more slowly and yielded lower ThT fluorescence amplitudes than in the presence of TCEP (Fig. 2e,f and Supplementary Fig. 2a,b; − vs +TCEP). Strikingly, C322S aggregated with kinetics comparable to reduced S320F_295-330_ but yielded 2-3 times higher ThT fluorescence amplitudes (Fig. 2e,f and Supplementary Fig. 2b,c; C322S vs +TCEP). TEM analysis revealed fibrillar assemblies under all conditions, although fibrils formed under non-reducing conditions appeared shorter and less ordered (Supplementary Fig. 3). Together, these data indicate that, in the context of the S320F_295-330_ peptide, non-reducing conditions slow aggregation kinetics while reducing conditions or mutation of the cysteine to serine accelerate fibril formation.

### C322S-containing tau S320F_295-330_ fibrils exhibit enhanced seeding in cells

Next, we tested whether different in vitro preparations of the S320F_295-330_ peptide, specifically reducing, or modifying C322, alter seed competency in cell-based tau aggregation assays. We used the tau P301S biosensor cell system^49^, which overexpresses tau repeat domain fused to FRET-compatible mClover3 and mCerulean3 fluorophores. Delivery of tau aggregates into biosensor cells via lipofectamine or through heparan proteoglycan-mediated macropinocytosis (“naked seeding”) converts diffusely distributed tau into aggregates, which are quantified by flow cytometry as the percentage of FRET-positive cells. For these experiments, larger batches of peptides were aggregated under non-reducing (no TCEP; (-)), reducing conditions (with TCEP; “red.”), and as the C322S mutant, with constant shaking at 37 °C for 3, 7, or 14 days (Fig. 3a). Fibril formation was confirmed by TEM (Supplementary Fig. 4). For seeding, fibrils were sonicated and delivered into cells by lipofectamine or naked seeding, followed by 3 or 4 days of growth, respectively (Fig. 3b). Cells were analyzed by flow cytometry following gating strategies to quantify the percentage of cells harboring FRET positive inclusions (Supplementary Fig. 5). In lipofectamine-mediated seeding assays, S320F_295-330_ C322S fibrils exhibited the highest activity, yielding 78% FRET-positive cells at 3 days, which declined to 53% and 31% at 7 and 14 days, respectively (Fig. 3c and Supplementary Fig. 6a). Seeding was highest at 60 nM and decreased at higher fibril concentrations (Fig. 3c and Supplementary Fig. 6a). S320F_295-330_ fibrils produced under non-reducing conditions exhibited maximum seeding of 40% at 60nM at day 3 which decreased to 10% by day 7, and 14 (Fig.3c and Supplementary Fig. 6a). S320F_295-330_ fibrils produced under reducing conditions showed similar but slightly lower signal to the non-reducing condition (20%, 6%, and 2% at 3, 7, and 14 days; Fig. 3c and Supplementary Fig. 6a). In naked seeding assays, we observed S320F_295-330_ C322S fibrils were the only condition to show robust and persistent activity (1-4%) across all incubation times and concentrations (Fig. 3d and Supplementary Fig. 6b). Reduced fibrils exhibited low seeding activity (1.3-2%; Fig. 3d and Supplementary Fig. 6b), and non-reduced fibrils produced <1% FRET-positive cells (Fig. 3d and Supplementary Fig. 6b). Notably, lipofectamine-mediated seeding declined with prolonged fibril aging, whereas naked seeding activity increased with additional in vitro incubation. Together, these results demonstrate that cysteine chemistry modulates the ability of this S320F_295-330_ sequence to form seed competent species. Mutation of C322 to serine had higher seeding activity compared to S320F_295-330_ fibrils produced in non-reduced and reduced S320F_295-330_ conditions. In naked seeding assays, only the C322S peptide fibrils retained activity.

**Figure 3.**
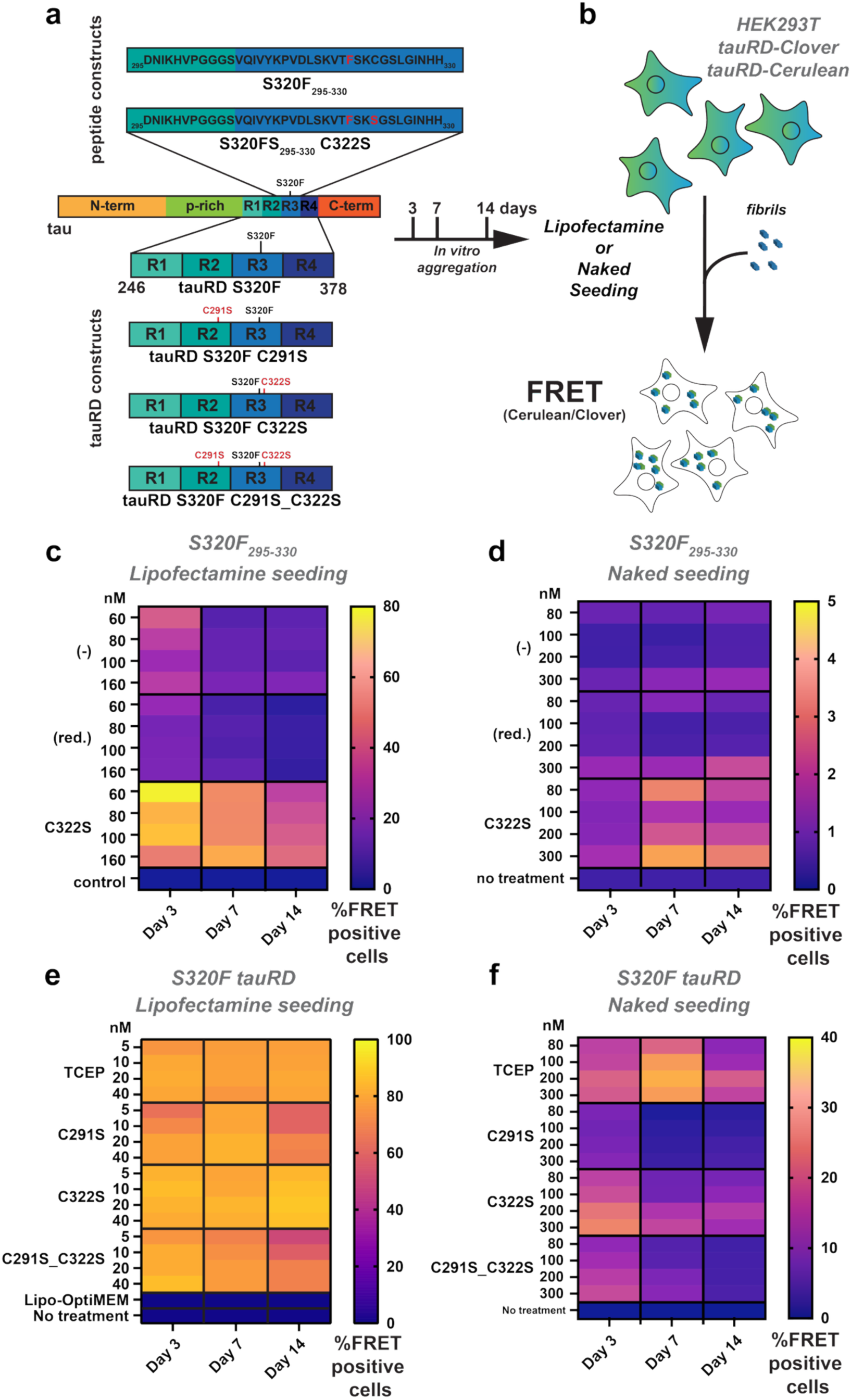
C322S mutation has different roles in tau fragment compared to tau repeat domain. (**a**) Schematic representation of S320F_295-330_ fragments and cysteine mutant (top) or tauRD S320F and cysteine mutants for the seeding experiment (bottom). Seeding experiments were carried out with the recombinant aggregate samples incubated in vitro for 3, 7 and 14 days and then delivered to cells using lipofectamine or directly (i.e. naked). (**b**) Schematic representation of cellular seeding workflow. Tau repeat domain spanning 246-380 (P301S tauRD) expressed as a C-terminal fusion to mCerulean3and mClover3. Peptide fibrils delivered using lipofectamine or directly (i.e. naked) and conversion of the intracellular tau into aggregates is measured using FRET on a flow cytometer. Percentage of FRET positivity was measured by flow cytometry with three technical replicates. (**c**) Percent FRET positivity of lipofectamine seeding is shown for S320F (-) (without TCEP), S320F (red.) and S320F C322S with four different concentrations (60, 80, 100 and 160 nM) delivered. End-point samples from day 3, 7 or 14 days of incubation used. Data shown as averages across as heat-maps as three technical replicates and colored from blue to yellow. (**d**) Percent FRET positivity of naked seeding of S320F (-) (without TCEP), S320F (red.) and S320F C322S with four different concentrations (80, 100, 200 and 300 nM). End-point samples from day 3, 7 or 14 days of incubation used. Data shown on heatmaps as averages across three replicates. (**e**) Percent FRET positivity from lipofectamine-mediated seeding activity of fibrils formed from tauRD S320F (TCEP), tauRD S320F C291S, tauRD S320F C322S and tauRD S320F C291S_C322S at 5, 10, 20 and 40 nM concentrations. Data shown as averages across three technical replicates and colored from blue to yellow. (**f**) Percent cell FRET positivity from naked seeding activity (in the absence of transfection reagents) of fibrils formed by incubating tauRD S320F (TCEP), tauRD S320F C291S, tauRD S320F C322S and tauRD S320F C291S_C322S preformed fibrils at 0, 80, 100, 200 and 300 nM with cells. All data is shown as the average of triplicates from 80nM to 300nM. Heat maps are shown in the plasma color scheme with purple and yellow denoting low and high percentage of cells with FRET positive aggregates, respectively.

### Cysteines in S320F tauRD in vitro fibrils contribute to seeding activity in cells

To test the role of the cysteine residues in the larger tau repeat domain (tauRD), we engineered tauRD with single and double mutants of C291 and C322 to serine encoding an S320F mutation (residues 246-378, herein tauRD): C291S, C322S and the double mutant C291S_C322S (Fig. 3a,e). The tauRD proteins were purified and aggregated in vitro at a concentration of 100 µM protein in 1x PBS (pH 7.4) in the presence of 5 mM TCEP and incubated for 3, 7 and 14 days (Fig. 3a). Formation of fibrils for each tauRD construct was confirmed using TEM at each time point (Supplementary Fig. 7a-d). Seeding activity was assessed at each time point in P301S tau biosensor cells^49^ using both lipofectamine-mediated and naked seeding assays (Fig. 3e). In the lipofectamine assay, all samples reached near-saturating seeding levels (∼80% FRET-positive cells at 40 nM) at 3 days. For tauRD S320F and tauRD S320F C322S, this activity remained stable through 7 and 14 days, whereas seeding by tauRD S320F C291S and the double cysteine mutant declined to ∼75% and ∼70% at later time points (Fig. 3e and Supplementary Fig. 8a). Consistent with this, seeding activity for tauRD S320F and tauRD S320F C322S was similar across all incubation times, while constructs containing the C291S mutation showed reduced activity after prolonged incubation (Fig. 3e and Supplementary Fig. 8a). Similar trends were observed in the naked seeding assay, where tauRD S320F and tauRD S320F C322S consistently produced higher FRET-positive signals than tauRD S320F C291S and the double cysteine mutant (Fig. 3f and Supplementary Fig. 8b). Notably, all constructs exhibited a time-dependent decline in naked seeding activity from 3 to 14 days. For S320F, seeding increased between 3 and 7 days before declining at 14 days, whereas all cysteine mutants displayed maximal activity at 3 days followed by progressive loss at later time points (Fig. 3f and Supplementary Fig. 8b). Together, these data indicate a complex role for the two cysteine residues in S320F-dependent tauRD aggregation in vitro and that cysteines may not be required to produce efficient seeds when delivered into the cytosol using lipofectamine but may play larger roles in in naked delivery where they must enter cells through oxidizing environments. By contrast, comparison with S320F_295-330_ peptide fibril experiments mutation of C322S yields seeds that are active via both delivery mechanisms.

### C322 but not C291 is required for S320F-based spontaneous aggregation

Next, we asked whether cysteine mutations alter the spontaneous aggregation phenotype of tauRD S320F previously observed in cells^28^. HEK293T cells were transduced with mEOS3.2-tagged tau repeat domain (tauRD, residues 246-380) constructs, including S320F and its cysteine mutants (C291S, C322S, and C291S_C322S). Control constructs included wild-type (WT) tauRD, the aggregation-prone P301S mutant, and β-sheet-disrupting proline substitutions in the essential ^275^VQIINK^280^ and ^306^VQIVYK^311^ amyloid motifs (I277P and I308P)^50^. mEOS3.2 is a photoconvertible fluorescent protein that forms a FRET-compatible green/red pair upon partial photoconversion. Spontaneous aggregation was monitored at 3, 5, 7, and 9 days post-transduction (Fig. 4a). At each time point, cells were harvested, photoconverted, and analyzed by flow cytometry to quantify the percentage of FRET-positive cells as a function of mEOS3.2 red fluorescence intensity (measured in arbitrary units 0-25k) (Fig. 4a and Supplementary Fig. 9a,b). At early time points (day 3), cells expressing tauRD S320F or tauRD S320F C291S exhibited low but detectable spontaneous aggregation (2-3%) at the highest expression levels (Supplementary Fig. 9b,c). By day 5, the FRET-positive population increased to 11.7% for S320F and continued to rise at days 7 and 9, reaching 36.2% and 72.6%, respectively (Fig. 4b and Supplementary Fig. 9b,c; 20k red intensity). TauRD S320F C291S followed a similar trajectory, with 2.7%, 7.6%, 17.4%, and 71.8% FRET-positive cells at days 3, 5, 7, and 9, respectively (Fig. 4b and Supplementary Fig. 9b,c). By contrast, tauRD S320F C322S remained near baseline through days 3-7 (0.36%, 0.51%, and 1.66%), increasing only modestly to 4% by day 9 (Fig. 4b and Supplementary Fig. 9b,c). TauRD S320F encoding the double cysteine mutant (C291S_C322S) showed no appreciable aggregation at any time point, remaining below 1% even at the highest expression levels and similar to WT tauRD, P301S tauRD and the aggregation-deficient tauRD I277P_I308P control (Fig. 4b and Supplementary Fig. 9b,c). The kinetics of aggregation of the tauRD S320F and tauRD S320F C291S in cells was reproducible in independent biological replicate experiments (Supplementary Fig. 9d,e). To corroborate these findings, cells were imaged at day 9 post-transduction. Constructs lacking FRET-positive signal by flow cytometry showed no visible inclusions by fluorescence microscopy imaged at day 9 (Fig. 4c and Supplementary Fig. 10-12). TauRD S320F and tauRD S320F C291S exhibited inclusion formation and cell-death at day 9 making the cell morphology more rounded in appearance and unsuitable for microscopy at this time point (data not shown). Despite lower FRET-positive signal tauRD S320F and tauRD S320F C291S show clear inclusion formation at day 3 (Fig. 4c and Supplementary Fig. 10). Together, these data indicate that C322 is critical for S320F-driven spontaneous tau aggregation in cells.

**Figure 4.**
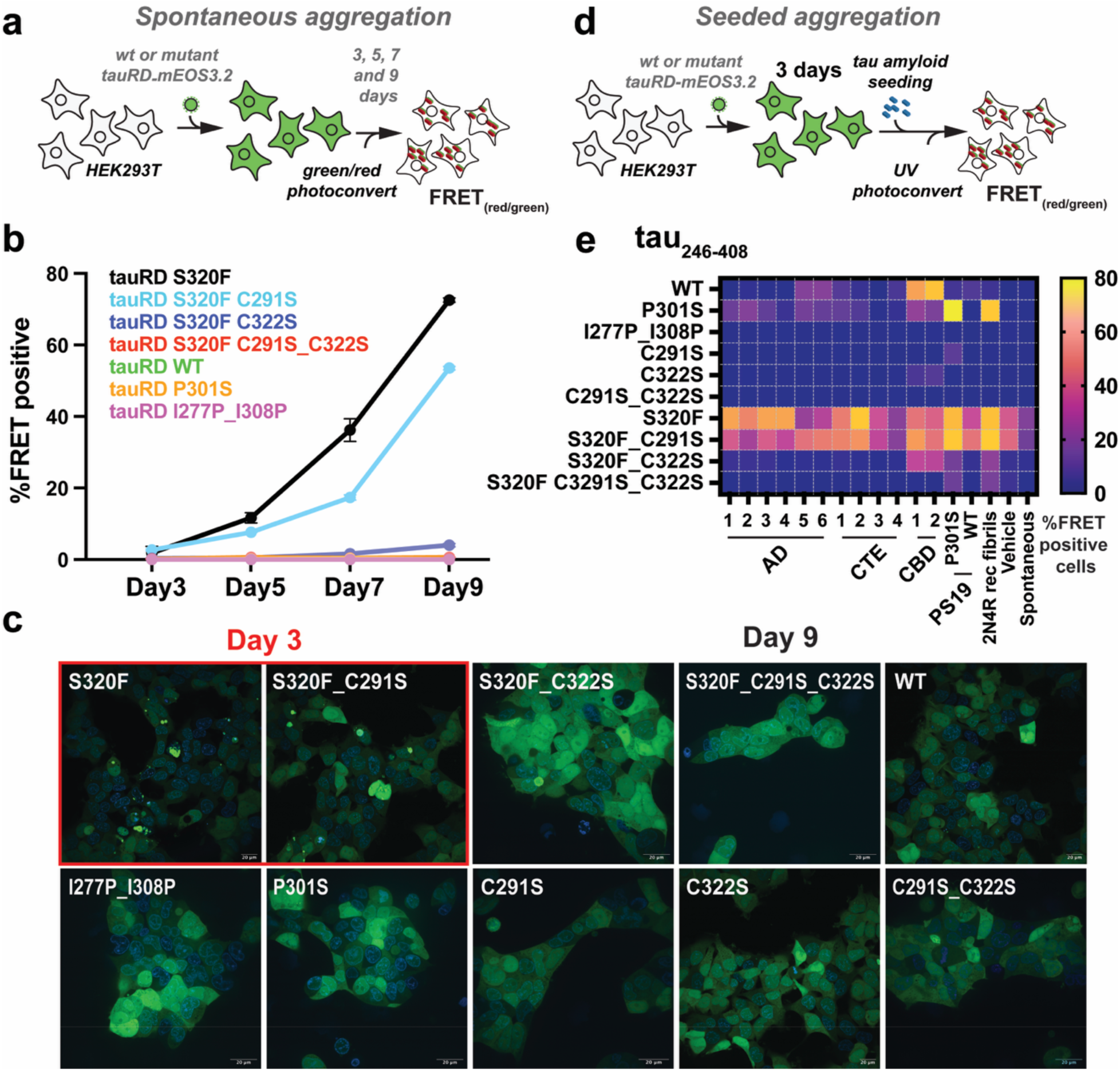
Cysteine-dependent modulation of spontaneous and seeded tau aggregation in cellular model system. (**a**) Schematic representation of spontaneous aggregation assay. HEK293T cell-line expressing WT or mutant tauRD-mEOS3.2 reporters were monitored over 3, 5, 7, and 9 days for aggregate formation, quantified by percentage of FRET positivity (PerCP-A FRET/mEOS3.2 Green) as a readout of intracellular tau assembly. (**b**) Quantification of spontaneous aggregation kinetics for WT tauRD and indicated disease-associated tauRD S320F and tauRD S320F cysteine mutant variants, plotted as percent FRET-positive cells over time (mean ± SEM.). (**c**) Representative fluorescence microscopy images showing intracellular aggregates at day 3 (left) and day 9 (right) for selected tauRD variants. Aggregates are visualized in the GFP channel with DAPI stained nucleus. Scale bar, 20µm. Constructs exhibiting robust early aggregation are highlighted. (**d**) Schematic of the seeded aggregation assay. HEK293T cell-line expressing WT or mutant tauRD-mEOS3.2 reporters were exposed to exogenous tau seeds delivered by lipofectamine, followed by assessment of intracellular aggregation after 3 days using FRET. (**e**) Heat map summarizing the percentage of FRET-positive cells for each tauRD variant under seeded conditions. Heat maps are shown in the plasma color scheme with purple and yellow denoting low and high percentage of cells with FRET positive aggregates, respectively.

We next used a centrifugation-based biochemical fractionation to quantify aggregate formation and assess whether disulfide-linked species could be detected. Lysates from cells expressing tauRD S320F and its cysteine mutants were collected at day 9, precleared, and ultracentrifuged at 186K to separate soluble species from large assemblies. Fractions were analyzed by SDS-PAGE and immunoblotting for tau (Supplementary Fig. 9f). The double cysteine mutant (C291S_C322S) showed the lowest amount of tau in the pellet fraction, whereas the highly aggregation-prone S320F construct did not yield the largest pellet signal. This indicates S320F aggregates might pellet during the lysate clarification step. Under non-reducing conditions, the cysteine-free C291S_C322S mutant lacked higher-molecular weight species and similar to S320F alone. By contrast, S320F C291S higher–molecular weight bands in both pellet and supernatant fractions, consistent with disulfide bond formation (Supplementary Fig. 9f). Interestingly, we also observed some evidence of higher-molecular weight bands for and S320F C322S at day 9, despite the low FRET signal (Supplementary Fig. 9f). These results suggest that disulfide bond formation is not the primary driver of tauRD S320F aggregation in cells, while highlighting that flow cytometry and biochemical fractionation may capture distinct aspects of aggregation behavior.

### Cysteine residues govern tau seeded aggregation across tauopathy seed types

To evaluate the contribution of cysteine residues to seeded tau aggregation, we compared seeding responses across a panel of tauRD constructs, including tauRD S320F and its cysteine mutants (C291S, C322S, and C291S_C322S), wild-type tauRD constructs (WT, C291S, C322S, and C291S_C322S), and established controls (P301S and the aggregation-deficient I277P_I308P mutant). All tauRD, tauRD S320F constructs were expressed with mEOS3.2 tag on the C-terminus and transduced into HEK293T cells. After 3 days, cells were seeded using lipofectamine-delivered aggregates derived from AD (n = 6), CTE (n = 4), CBD (n = 2) patient tissues, PS19 tauopathy and control mouse brains^51^ (P301S_PS19 and WT_PS19), or recombinant heparin-assembled 2N4R tau. Aggregation was quantified 3 days later by partial photoconversion of mEOS3.2 followed by flow cytometry to measure the percentage of FRET-positive cells (Fig. 4d and Supplementary Fig. 9a). TauRD S320F and S320F C291S showed the highest raw FRET signals (Fig. 4e; 15k mEOS3.2_red_), but these measurements were confounded by spontaneous aggregation. As expected, P301S tauRD responded strongly to PS19 lysates (77%) and recombinant fibrils (68%), whereas WT tauRD showed preferential sensitivity to CBD-derived seeds (59-68%) with minimal response to other seed types (Fig. 4e and Supplementary Fig. 12). The tauRD dual proline mutant (I277P_I308P) was insensitive to all seeds, yielding near-background signal (0.13-0.7%), consistent with its inability to be seeded (Fig. 4e and Supplementary Fig. 11). Strikingly, introduction of cysteine mutations into WT tauRD markedly reduced seeding across all seed types. TauRD C291S responded weakly only to PS19 and recombinant fibrils, while tauRD C322S showed modest responses exclusively to CBD-derived seeds (Fig. 4e and Supplementary Fig. 12). The tauRD double cysteine mutant (C291S_C322S) was essentially insensitive to all seeds, phenocopying the dual proline mutant (Fig. 4e and Supplementary Fig. 12). Analysis of seeding responses for the S320F variants showed that tauRD S320F and tauRD S320F C291S behaved similarly across seed types (Fig. 4e). By contrast, tauRD S320F C322S responded selectively to CBD-derived seeds, PS19 lysates, and recombinant fibrils, with minimal response to other tauopathy seeds (Fig. 4e and Supplementary Fig. 10). The tauRD S320F C291S_C322S double mutant was largely resistant to seeding to all seeds except PS19 and recombinant fibrils (Fig. 4e and Supplementary Fig. 10). Together, these data indicate that C322 is required for spontaneous aggregation of tauRD S320F and that combined mutation of C291 and C322 renders both WT tauRD and tauRD S320F insensitive to seeded aggregation. Notably, the differential effects of C291S and C322S across seed types, particularly the preserved response to CBD-derived seeds, may suggest that the two cysteines play distinct, strain-dependent roles in tau seeding.

### Cysteines rank with amyloid motifs as essential regulators of seeded tau aggregation

We and others have previously shown that mutation of essential residues within the ^275^VQIINK^280^ and ^306^VQIVYK^311^ amyloid motifs to proline abolishes tau aggregation, underscoring the central role of these elements in tau assembly^19–22,24,50^. Our cell-based data further indicate that cysteine-to-serine substitutions profoundly affect both S320F-driven spontaneous aggregation and seeded tau assembly. To place the contribution of cysteine residues in the context of an extended tauRD sequence spanning residues 246-408, we performed a systematic alanine-scanning mutagenesis across tauRD^52^ and quantified the effect of each mutation on seeded aggregation using lysates derived from tauopathy patient brain samples, including AD (n = 4), CBD (n = 4), and PSP (n = 4). HEK293 cells were transduced with lentivirus encoding each tauRD alanine mutant (residues 246-408) fused to mEOS3.2, seeded with lipofectamine-delivered tissue-derived lysates, and analyzed after 3 days by flow cytometry following partial photoconversion of mEOS3.2 (Supplementary Fig. 14). Mutants were tested in duplicate for each tissue, and mean values were converted to Z-scores to normalize across the mutant series and account for intrinsic differences in seeding efficiency. Average Z-scores were then compared across AD, CBD, and PSP samples (Fig. 5b; AD in blue, CBD in red, PSP in green).

**Figure 5.**
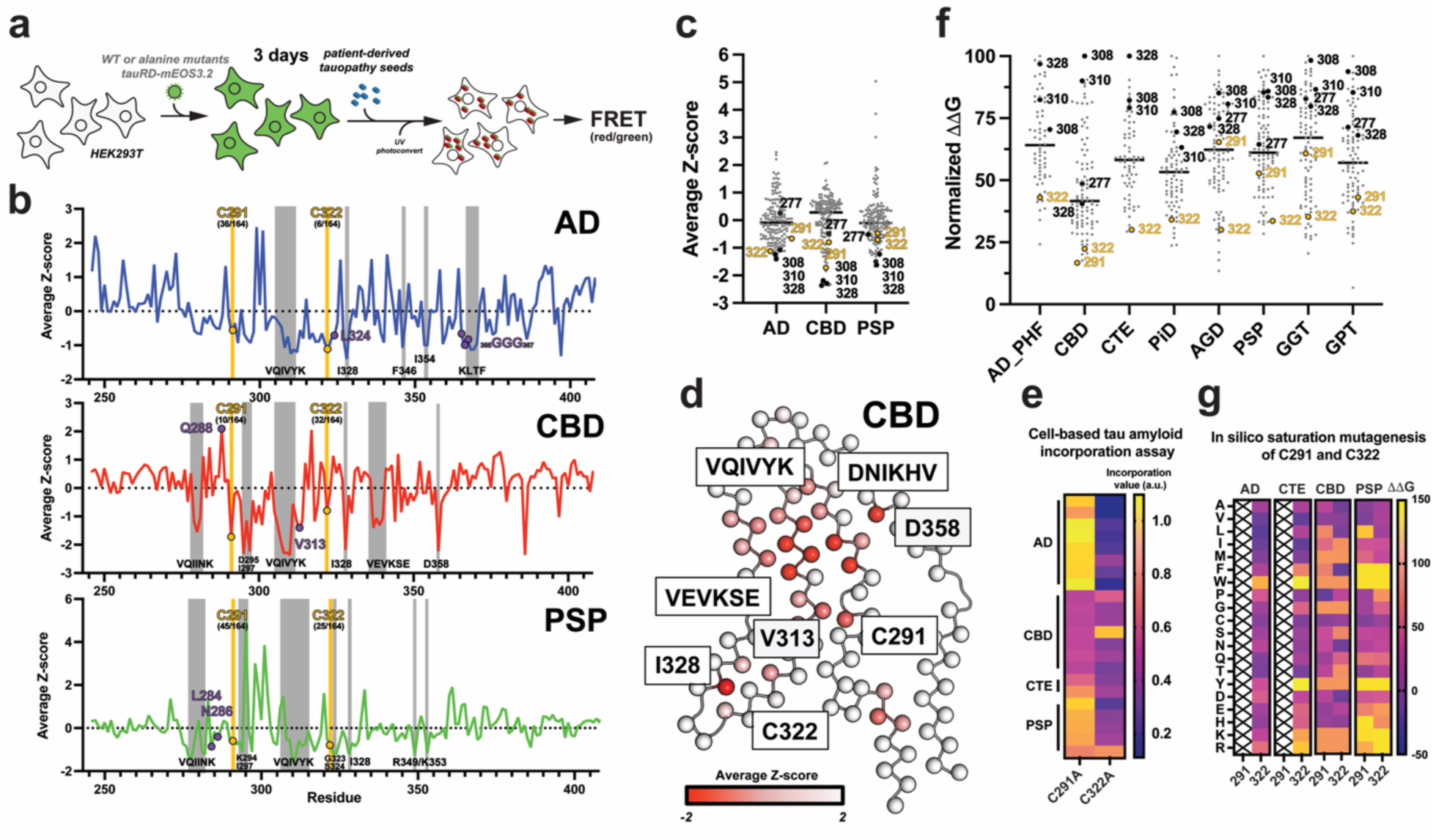
Disease-specific tau sequence features and energetic hotspots linked to aggregation and seeding. (**a**) Schematic of the alanine scan seeding cellular aggregation assay. HEK293T cell-line expressing tauRD-mEOS3.2 reporters are monitored for spontaneous aggregation or following exogenous seeding with patient material lysates, with intracellular assembly quantified by FRET as a readout of tau-tau interactions. (**b**) Z-scores from seeding alanine scan mapped along the tau repeat domain for Alzheimer’s disease (AD), corticobasal degeneration (CBD), and progressive supranuclear palsy (PSP) colored blue, red and green, respectively. Traces highlight regions of elevated aggregation propensity, with key amyloidogenic motifs (including VQIVYK) and cysteine-containing segments marked. Gray bars indicate regions enriched in disease-associated energetic contributions. Yellow points highlight location of cysteine 291 and 322. Purple positions indicated amino acids that form interactions with the cysteine residues. (**c**) Distribution of residue-level Z-scores from each alanine mutant seeded with AD, CBD, and PSP tissues. Data is shown as grey points with hotspot residues colored black and cysteines colored yellow. (**d**) Structural mapping of the Z-score residue level derived from the alanine scan seeding experiment mapped onto CBD tauopathy fibril structure. Residues and motifs that have low Z-score values are labeled. Structures of fibril layer are shown as wire frame with C-β atoms shown as spheres. Values are colored as a gradient from white (positive Z-score) to red (negative Z-score). (**e**) Heat map illustrating efficiency of tauRD alanine mutant incorporation into preformed AD, CBD CTE and PSP aggregates as measured as a fraction of incorporation. Heat maps are shown in the plasma color scheme with purple and yellow denoting low and high incorporation, respectively. (**f**) Residue-level ΔΔG values calculated by comparing WT and alanine residues for each position in AD, CBD, PSP, CTE, PiD, AGD, GGT and GPT tauopathy fibril structures. Data is shown as grey points with hotspot residues colored black and cysteines colored yellow. (**g**) Heat map summarizing predicted changes in ΔΔG of fibrils upon mutation of cysteine 291 and 322 systematically to each of the 20 amino acids. Heat maps are shown in the plasma color scheme with purple and yellow denoting low and high ΔΔG values, respectively.

As expected, alanine substitutions within established amyloidogenic motifs, including ^306^VQIVYK^311^, ^275^VQIINK^280^, and ^335^VEVKSE^340^, strongly inhibited seeded aggregation across all tauopathies (Fig. 5b, gray bars). Additional nonpolar residues previously implicated in fibril stability, such as ^375^KLTF^378^, I297, I328, and F346^22,24^, also ranked among the strongest aggregation suppressors. Strikingly, C322A emerged as one of the most potent inhibitors of AD-seeded aggregation (ranked 6 out of 164), with an effect only comparable to core amyloid residues, whereas C291A had a more modest effect (ranked 36 out of 164) (Fig. 5b, top panel). By contrast, CBD-seeded aggregation was strongly inhibited by C291A (ranked 10 out of 164) but only weakly affected by C322A (ranked 32 out of 164) (Fig. 5b, middle panel). For PSP, where signal-to-noise was lower, both C291A and C322A nonetheless reduced seeded aggregation (ranked 45 out of 164 and 25 out of 164, respectively) (Fig. 5b, bottom panel). Consistent with these rankings, cysteine-to-alanine mutants (Fig. 5c, yellow points) clustered with known aggregation-critical positions in the Z-score distribution (Fig. 5c, black points), placing them among the most crucial residues governing seeded tau aggregation (Fig. 5c, black points). Mapping these effects onto tauopathy-specific fibril structures revealed that essential residues localize to networks of stabilizing interactions, particularly in CBD fibrils where they converge around the ^306^VQIVYK^311^ amyloid motif (Fig. 5d and Supplementary Fig. 15a) consistent with prior in silico thermodynamic profiling^22^.

To further assess the role of cysteines in fibril propagation, we reanalyzed a previously published assay that quantified incorporation of mutant tau sequences into pre-existing tauopathy-specific aggregates derived from patient brain samples. In this assay, disease-derived aggregates were first used to template conversion of wild-type tauRD, followed by transduction of alanine-substituted tau variants to assess required determinants of incorporation into existing aggregates^52^. Focusing on C291A and C322A, we found that C322A markedly reduced incorporation into AD, CTE, and PSP aggregates, whereas C291A had little to no effect; by contrast, both mutations only modestly affected incorporation into CBD aggregates (Fig. 5e). These results mirror the seeding data (Fig. 5a,b) and demonstrate that cysteine residues regulate not only seeded aggregation but also tau incorporation into pre-existing fibrils in a tauopathy-dependent manner. It is thus possible that the cysteine residues may stabilize interaction of tau monomers with seeds or growing aggregates to promote efficient elongation.

To gain mechanistic insight into how cysteines influence tau aggregation, we reexamined previously published thermodynamic ΔΔG calculations comparing wild-type and alanine-substituted tau amyloid assemblies^22^. These analyses identified core amyloid motifs ^306^VQIVYK^311^, ^275^VQIINK^280^, and ^335^VEVKSE^340^) and the PAM4 motif (^349^RVQSKIG^355^) as major contributors to fibril stability^22,25^. Reanalysis revealed that, unlike the alanine scan seeding data, cysteine residues do not rank among the most stabilizing positions based on ΔΔG predictions (Fig. 5f). Consistent with this, structural analysis shows that cysteines are buried within fibril cores and primarily engage in nonpolar interactions with neighboring residues or adjacent strands (Supplementary Fig. 16). In AD and CTE fibrils, C322 contacts residues such as ^365^GGG^367^ and L324, whereas in the more complex CBD and PSP folds, cysteines interact with nonpolar side chains and aliphatic portions of asparagine or glutamine residues (Supplementary Fig. 16). Importantly, mutation of residues contacting cysteines produced highly variable effects in the seeding assay and none recapitulated the magnitude of the cysteine-to-alanine mutations (Fig. 5b, purple dots; Supplementary Fig. 15b). Similarly, ΔΔG predictions for these contacting residues spanned a broad range, with some mutations exerting minimal effects and others contributing partially to fibril stability (Supplementary Fig. 15c). Finally, we assessed tolerance at C291 and C322 by in silico mutagenesis across AD, CTE, CBD, and PSP tau folds, estimating the ΔΔG for substitution of cysteine with all amino acids. We find that on average large nonpolar (i.e. trp) or charged residues (i.e. asp/glu and arg/lys) negatively impact stability of fibrils (Fig. 5g). By contrast, smaller amino acids and in AD/CTE even larger nonpolar residues can be accommodated in place of the cysteine residues (Fig. 5g). This data suggest that cysteine residues appear to not be optimally packed in these structures suggesting that their packing may not be their defining contribution. Together, these data support a model in which cysteine residues play a disproportionately larger role in promoting seeded tau aggregation through mechanisms that are not captured by thermodynamic stability alone, with C291 and C322 exerting distinct, tauopathy-dependent effects.

## Discussion

The FTD-associated S320F mutation is notably efficient in its ability to drive cell-autonomous, spontaneous aggregation of both full-length tau and tau fragments in vitro and in cells, distinguishing it from other disease-linked MAPT mutations and other tau variants^28,42,43^. Our data demonstrate that S320F directly participates in the fibril core and reshapes the local chemical environment to favor amyloid formation (Fig. 6). In particular, burial of the phenylalanine side chain alters hydrophobic packing that stabilize dimers and facilitating intermolecular disulfide formation. Consistent with this model, recent structures of tau fibrils isolated from animals expressing full-length tau that combined two FTD-linked mutations, P301L and S320F, revealed a critical role for the phenylalanine substitution in stabilizing the fibril core, while also adopting a more collapsed architecture likely driven by the additional P301L mutation, which perturbs interactions surrounding the essential ^306^VQIVYK^311^ amyloid motif^43^. In FTD-tau patients harboring the S320F mutation, the phenylalanine forms stabilizing interactions with I328 and L325^44^. In our structure, the stability is conferred by nonpolar interactions between ^306^VQIVYK^311^ amyloid motif interactions and hydrophobic residues near the C-terminus reminiscent of early fibrillar intermediates that are on path toward mature fibrils^20,27^. Our structure shows the C322-C322 stabilizing parallel dimer interfaces; however, in solution, alternative antiparallel geometries may also be accessible. Reducing conditions likely relax constraints imposed by disulfide-mediated dimerization, allowing peptide-peptide interactions to dictate productive folding, followed by oxidation in the parallel orientation into this fibril state and may explain the higher aggregation propensity of the C322S mutant. Surprisingly, the S320F side-chain itself contributes local overpacking rather than direct energetic stabilization which is likely compensated by the intermolecular C322-C322 disulfide bond linking adjacent chains. To our knowledge, this represents the first tau fibril structure in which an intermolecular disulfide directly stabilizes interchain contacts. Together, these findings provide a mechanistic framework for how a single nonpolar substitution can rewire the conformational ensemble of tau to favor self-assembly, explaining the aggressive, seed-independent aggregation phenotype associated with S320F.

**Figure 6.**
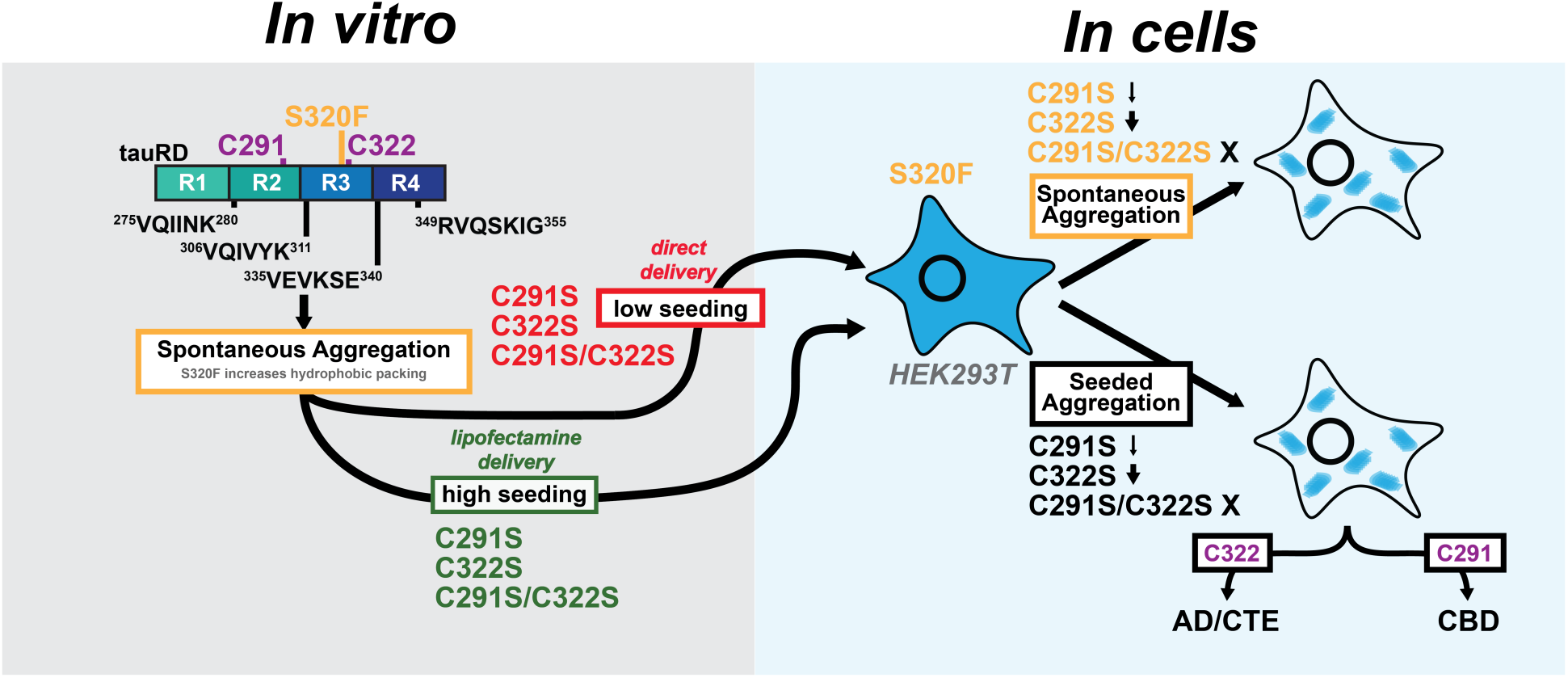
C291 and C322 act as chemical gatekeepers that determine whether tau forms productive, strain-competent assemblies or off-pathway aggregates. (Left) The S320F mutation introduces a local destabilizing, hydrophobic residue that exposes amyloidogenic motifs and allows for increased hydrophobic packing, leading to spontaneous aggregation of tauRD. Cysteine residues are not required for fibril formation in the context of S320F tauRD to enable seeding using lipofectamine, however, aggregates with cysteines appear to preserve more seeding when delivered into cells directly. (Right) Perturbations at flanking cysteine residues decrease spontaneous aggregation and strain-dependent seeded aggregation with the double C291S and C322S mutant inhibiting both.

Functional assays further reveal that the contribution of cysteine residues to tau aggregation is highly context-dependent. While in vitro-assembled S320F_295-330_ fibrils seed efficiently even in the absence of C322, tau repeat domain constructs encoding S320F require intact cysteines to retain seeding activity following direct cellular delivery (Fig. 6). Prior studies of the dGAE tau fragment that encodes residues 297-391 similarly reported enhanced aggregation and seeding upon C322 mutation, suggesting that constructs containing a single cysteine behave differently from those encoding both C291 and C322^53^. Structural studies of C322S-containing dGAE tau fragments further showed increased recombinant yields and adoption of folds partially resembling Alzheimer’s disease tau strains^54^. Together, these observations point to a complex coupling between the two cysteine positions that cannot be inferred from studies of isolated cysteine substitutions alone.

Our data also provide compelling evidence that cysteine residues play a central role in tau amyloid formation in the reducing environment of cells. The C291 and C322 positions embedded within highly conserved SKXGS motifs^55^ in each of the four tau repeat domains that stabilize tau-microtubule interactions^23,56^. Post-translational modification within these motifs has been shown to regulate tau assembly^57^ and inhibit microtubule binding^58^. Early work by Mandelkow and colleagues suggested that intramolecular C291-C322 disulfides in tau inhibit aggregation, whereas intermolecular C322-C322 disulfides may promote tau assembly in vitro, leading to the widespread use of reducing conditions in tau aggregation assays. More recent work demonstrated a seeding barrier between oxidized and reduced tau fibrils^59^, indicating that redox state can encode distinct fibril conformations. However, the functional relevance of cysteine chemistry in physiological aggregation contexts has remained largely unexplored, with only limited studies showing reduced tau toxicity upon cysteine mutation in *Drosophila* models^60,61^.

Our study is the first to systematically interrogate the role of cysteines in cellular models of both spontaneous and seeded tau aggregation. We show that cysteine residues are essential for S320F-driven spontaneous aggregation and, critically, play strain-specific roles in seeded aggregation of wild-type tau (Fig. 6). These findings are consistent with uptake studies showing that recombinant tau seeds become oxidized upon cellular entry^62^, suggesting that oxidizing environments encountered by the seed during uptake may preserve or enhance cysteine-dependent seed competence^63^. Interestingly, in a prior study we showed that covalent crosslinking model peptide fragments into dimers that encode the ^306^VQIVYK^311^ motif yielded species that efficiently self-assembled and seeded wild-type tau^64^. These data provide a potential explanation for why cysteine-containing seeds are more effective at physiological propagation and may also rationalize differences observed between single cysteine peptide-based and two cysteine containing tauRD-based seeds. Our alanine mutagenesis seeding screen reinforces this conclusion, identifying C291 and C322 among the most influential residues governing strain-dependent cellular seeding, ranking comparably to core amyloid residues. This underscores the idea that cysteines encode unique chemical functionality that cannot be readily substituted and highlights a potential role for cellular redox state in modulating tau propagation. Notably, C291 and C322 play distinct roles in the S320F background: one appears to promote productive aggregation while the other is dispensable. This also extends to seeded aggregation, where AD-patient derived seeds require C322, while CBD-derived patient seeds are more dependent on C291. This suggests that the cysteines play sequence and structure context-dependent roles in tau assembly. While our study focused on tau assembly, recent studies on TDP-43 showed that cysteine residues in an RNA binding RRM motif contribute to its pathogenic misfolding and that mutation of these cysteines prevents formation TDP-43 assemblies^65^. Thus, it is possible that tuning intermolecular disulfide bond formation in amyloidogenic proteins through changing redox potential or using modifications may be a plausible strategy to suppress pathogenic seed formation beyond tau.

Together, our findings support the prion hypothesis by demonstrating how protein structure and chemistry encode disease-relevant behavior. The S320F mutation exemplifies how a single amino acid substitution can generate a tau conformer capable of autonomous aggregation, bypassing the need for templated aggregation. More broadly, our results identify cysteine residues as chemically tunable control points in tau assembly and propagation. Targeted modulation of cysteine chemistry, through redox control or covalent intervention, may offer new opportunities to attenuate strain-specific tau propagation. Future studies must directly test how disulfide formation biases tau toward pro- or anti-aggregation states in cells and in vivo. The outsized role of cysteine residues uncovered here reframes tau aggregation as a chemically regulated process and opens new avenues for diagnosis and therapeutic intervention in tauopathies.

## Materials and Methods

AD, CBD, and PSP human brain tissue was obtained through the University of Texas Southwestern Medical Center Neuropathology Brain Bank. Legally binding informed consent for research use of postmortem brain tissue was obtained from each decedent’s next of kin in accordance with Texas law. Since all tissue was obtained from nonliving subjects, such research was considered exempt from human subject regulations by the IRB. Select AD, PSP and CDB cases were obtained from the Alzheimer’s Disease Research Center (ADRC) from Washington University in St. Louis and CTE cases were obtained from the CTE Center at Boston University. A summary of cases is detailed in Supplementary Table 4.

### Peptide synthesis

S320F_295-330_ and cysteine mutants tau peptide fragments (C322S), were synthesized with N-terminal acetylation and C-terminal amidation and purified to >95% by Genscript.

### Aggregation of S320F tau_295-330_ and variant peptides

Peptides were disaggregated using trifluoroacetic acid (TFA, Sigma-Aldrich) at a ratio of 2 mg of peptide/200 mL of TFA and kept overnight in a fume hood to remove the organic solvent. Any residual solvent was removed by purging N_2_ gas, followed by brief resuspension in 200 mL MilliQ water prior to being flash frozen in liquid nitrogen and lyophilization. Disaggregated peptide powder was stored in -20°C until needed. The disaggregated peptide was dissolved in 1x phosphate-buffered saline (PBS) (136.5 mM NaCl, 2.7 mM KCl, 10 mM Na_2_HPO_4_, 1.8 mM KH_2_PO_4_, to the desired concentration. For kinetic experiments, S320F tau_295-330_ and variant peptides were incubated at a range of concentrations between 10, 15 and 20 uM. For kinetic experiments, Thioflavin-T (ThT, Sigma-Aldrich) was added to a final concentration of 25 mM. For reducing conditions, 5 mM tris(2-carboxyethyl) phosphine hydrochloride (TCEP Sigma-Aldrich). 55 mL of the sample was added (in triplicate) to wells in a 384-well clear-bottom plate (Corning). Kinetic aggregation experiments were carried out using a Tecan Spark plate reader. The measurements were recorded at an excitation of 446 nm and emission of 486 nm, with 10 seconds shaking prior to each read, with an interval of 30 minutes. The data points were plotted as averages with standard deviation, and t_1/2_ for each experiment was determined using a non-linear sigmoid fit in GraphPad Prism v10.0. For cell-based seeding experiments, S320F tau_295-330_ and variant peptides were aggregated at 200 µM in 1x PBS (pH 7.4) in a 200 µl volume. For reducing conditions, 5 mM tris(2-carboxyethyl) phosphine hydrochloride (TCEP Sigma-Aldrich) was added. For non-reducing conditions, TCEP was not included. The samples were incubated at 37 °C with interval mixing (15 seconds on at 800 rpm, 10 minutes off) on a Thermomixer (Eppendorf) for 3, 7, and 14 days.

### Transmission electron microscopy

A 5 μL aliquot of the aggregated fibril sample was placed on a glow-discharged 300 mesh copper grid for 2 min and negatively stained with 5 μL of 2% uranyl acetate for 45 sec. The grid was washed twice for 30 s with MilliQ water to remove excess staining solution and dried. Images were acquired in the Electron Microscopy Core facility at UT Southwestern Medical Center using a Tecnai G2 Spirit and a JEOL 1400 Plus electron microscopes operating at 120 kV.

### Cryo-electron microscopy data collection

The filaments were assembled by incubating the peptide at a concentration of 300 µM in the presence of 2 mM TCEP in 1x PBS (pH 7.4) at 37 °C with interval mixing (15 seconds on, 10 minutes off) on a Thermomixer (Eppendorf) for 72 hours. 3.5 μL of this reaction was applied to glow-discharged Quantifoil R1.2/1.3, 300 mesh copper grids that were plunge-frozen in liquid ethane using a Vitrobot with the following settings: blotting force 0, blotting time 30 s, humidity 98-100 % at 4 °C. Cryo-EM data were acquired using a FEI Titan Krios 300 kV transmission electron microscope. Images were recorded with a total dose of 60 electrons per Å^2^ and a pixel size of 0.8334 Å. A total of 6,509 movies were collected with 105,000x magnification. See Table 1 in the supplementary information for further details.

### Helical reconstruction of fibrils

The dataset was processed and analyzed using helical reconstruction in RELION 4.0 ^45^. The raw movie frames were gain-corrected, motion-corrected and dose-weighted using RELION’s own implemented motion correction software. Contrast transfer function (CTF) estimation was carried out using CTFFIND 4.1^66^. Particles from 6,509 micrographs were picked manually, and segments were initially extracted with a box size of 1,024 pixels to facilitate crossover distance determination. In parallel, an independent set of segments was extracted using a smaller box size of 384 pixels and downscaled to 128 pixels for multiple rounds of two-dimensional (2D) classification. The best looking 2D classes were selected and re-extracted at the original pixel size with a box size of 384 pixels, followed by an additional round of 2D classification. The highest estimated resolution 2D classes that spanned the crossover were selected to generate initial three-dimensional (3D) model using relion_helix_inimodel2d script. Refine3D were performed (reference model was low-pass filtered to 10 Å), followed by a 3D classification into 4 classes without image alignment option. The best-looking classes were selected and subjected to further Refine3D rounds for helical parameter optimization. Multiple rounds of Bayesian polishing, Refine3D and CTF refinement were used to improve map quality and resolution. The final map was sharpened using RELION postprocessing procedures and resolution estimates were calculated based on the Fourier shell correlation (FSC) between two independently refined half-maps at 0.143.

### Model building and refinement

The refined map was further sharpened using phenix.auto_sharpen at the cutoff resolution of 3.7 Å^67^. Multiple rounds of model refinement were carried out using the default settings in phenix.real_space_refine with non-crystallographic symmetry constraints, minimization_global, rigid_body, and local_grid_search^68^. Model geometry was evaluated using MolProbity, a built-in tool in Phenix^69^. After each round of refinement, problematic or poorly fitting regions in the model were manually modified using COOT^46^. This procedure was repeated until we achieved an acceptable stereochemistry model and high overall correlation coefficients of model:map.

### *In Silico* Rosetta-based thermodynamic profiling of S320F_295-330_ and tauopathy fibril structures

3-, 6-, 9- and 12-layer assemblies of the cryo-EM structure were prepared using an in-house script. These assemblies were then used as input for the subsequent mutagenesis and minimization using the RosettaScripts interface to Rosetta^70^ in framework similar to a previously described method^22^. Changes in assembly energy were calculated using a method adapted from the Flex ddG^71^ for amyloid fibrils^22^. For the 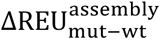 calculations, all chains were mutated. From the input assembly, a set of pairwise atom constraints with a maximum distance of 9 Å were generated with a weight of 1, using the fa_talaris2014 score function. Using this constrained score function, the structure then underwent minimization. After minimization, the residues within 8 Å of the mutation site underwent backrub sampling to better capture backbone conformational variation. These sampled structures were either only repacked and minimized, or the alanine mutation was introduced, followed by repacking and minimization. For 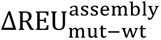 calculations, the bound wild-type and bound mutant structures reported by the interface ddG mover were used for estimating the change in assembly energy due to an alanine substitution. This is repeated for 35 independent replicates. The lowest energy bound mutant and bound wild-type structure energies from each replicate were extracted, and the change in energy as calculated by subtracting the wild-type, non-mutagenized assemblies’ energy from the mutant assemblies’ energy. This procedure is repeated for every residue in the structure to generate a set of 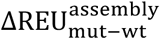 values. This method repeated using tauopathy fibril structures to generate a set of 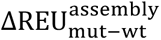 values for each unique fibril structure. The adapted Flex ddG protocol was further used to perform saturation mutagenesis at positions C291 and C322 across tauopathy fibril folds associated with AD (PDB ID: 6HRE), CTE (PDB ID: 6NWP), CBD (PDB ID: 6TJO), and PSP (PDB ID: 7P65). Saturation mutagenesis followed the same workflow described above for alanine scanning but expanded to include substitution of cysteine residues at positions 291 and 322 with all 20 amino acids. For each mutation, structures were locally minimized and evaluated across 35 independent replicates. Reported values represent the mean energetic change across replicates, with assembly stability quantified as 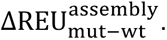

### Expression, purification and aggregation of tauRDS320F and cysteine mutants

TauRD S320F and cysteine mutants (C291S, C322S and C291S_C322S) were cloned into the pet29b (+) backbone using gene blocks from IDT. Plasmids were transformed into BL21(DE3), and single colonies were inoculated into 20 mL LB media with 20 µg/ml Kan and incubated overnight at 37 °C. Overnight culture was diluted to 1L LB with 20 µg/ml Kan and incubated at 37 °C with 220 rpm shaking. The protein expression was induced with 1 mM isopropyl β-D-1-thiogalactopyranoside for 4 h at 37 °C. The cells were harvested by centrifuging the cultures at 4000 × *g* for 20 minutes and lysed in 20 mM 2-(*N*-morpholino) ethanesulfonic acid (MES) pH 6.8, 1 mM Ethylenediaminetetraacetic acid (EDTA), 1 mM MgCl_2_, 5 mM β-mercaptoethanol. The lysate was sonicated using a probe sonicator at 40-50 % power, 5x pulse, 15 minutes with on/off cycles.

The lysate was centrifuged, NaCl was added to a final concentration of 500 mM, and the supernatant was boiled for 20 min in a water bath. The lysate was then centrifuged at 15,000 × *g* for 20 minutes. The supernatant collected after centrifugation was dialyzed against 20 mM MES, 50 mM NaCl, 5 mM β-mercaptoethanol, pH 6.8. The lysate was filtered using a 0.22 µm filter, loaded on a 5 ml HiTrap SP HP cation exchange column, and eluted with NaCl gradient (50 mM–800 mM) in 20 mM MES, 50 mM NaCl, 5 mM β-mercaptoethanol, pH 6.8. The eluted fractions were concentrated using a 10 kDa cutoff Amicon concentrator (EMD Millipore) and buffer exchanged into 1x PBS (pH 7.4) containing 5 mM β-mercaptoethanol and stored at - 80°C. In vitro aggregation experiments on tauRD S320F and cysteine variants were performed at a concentration of 100 µM in 1x PBS (pH 7.4) in the presence of 5 mM TCEP (in a 200 µl volume. The samples were incubated at 37 °C with interval mixing (15 seconds on at 800 rpm, 10 minutes off) on a Thermomixer (Eppendorf) for 3, 7 and 14 days.

### Cell-based seeding assay with S320F_295-330_ peptide fragment and *in vitro* aggregated tauRD S320F and tauRD S320F cysteine mutants

The NK13 biosensors cell lines established in the Diamond lab (ABM, cat. T3488) expressing tauRD (246-380) with P301S mutation fused to mCerulean3 or mClover3 were plated in a 96-well plate at 10,000 cells per well in 200 ml of complete DMEM media with 10% FBS and treated after 18 hours of plating. The fibrils were diluted to a working concentration of 1 µM and sonicated at an amplitude of 65 for 30 sec on and 30 sec off with a total on time of 2 min using a water bath sonicator prior to seeding on cells. For lipofectamine seeding experiments, 0.75 ml of Lipofectamine 2000 (Invitrogen) and 9.3 mL Opti-MEM (Gibco) for each well of a 96-well plate was incubated as a master-mix for 5 min prior to being added to sonicated 295-330 peptide/tauRD fibril. Opti-MEM was added to reach a final volume of 260 ml and a desired concentration of peptide/tauRD fibril (60 nM to 160 nM). The mixture was incubated for 20 min at RT. For seeding with tauRD sequences, the concentration ranged from 5 nM to 40 nM. Each concentration was performed as a biological replicate. For naked seeding experiments, 5000 cells/well were plated in 100 ml of complete media in a 96-well plate. Sonicated fibrils were diluted in the DMEM complete media (Gibco) with 10% FBS (Corning) and added to the cells dropwise with final concentrations ranging from 80 nM to 300 nM for both peptide and tauRD sequences. The cells were harvested after 96 hours of treatment by 0.25% trypsin digestion and fixed in 1x PBS. Cells were harvested after 72 hours of lipofectamine seeding and 96 hours of naked seeding by 0.25% trypsin and then fixed in 1x PBS with 2% paraformaldehyde (Electron Microscopy Sciences #15710). Fixed cells were resuspended in 1x PBS and were run on flow cytometer to quantify the % FRET positivity.

### Spontaneous aggregation assay and seeding assay for tauRD mEO3.2 constructs

TauRD constructs (residues 246-380) that are C-terminal fused to mEOS3.2 were tested for spontaneous aggregation in parental HEK293T cell line (ATCCCRL-3216). The following mEOS3.2 constructs: tauRD S320F, tauRD S320F C291S, tauRD S320F C322S, tauRD P301S, tauRD S320F C291S_C322S (cysteine double mutants), tauRD I277P I308P, tauRD C291S, tauRD C322S, tauRD C291S_C322S were cloned in a FM5 lentiviral vector backbone described previously in Vaquer-Alicea et al. 2025. All tauRD constructs were cloned with with the minimal tau repeat domain, residues 246-380. The WT tauRD fragment was cloned with a C-terminal extension (residues 246-408) that increases seeding responses of WT tau. For the lentivirus production 300,000 cells were plated in each well of a six-well plate which were 18 hours later transfected with mEOS3.2 plasmids at 1.6 mg with 1.2 mg of PSP-G and 0.4 mg of VSV-G helper plasmids using 7.5 ml of TransIT-293 (Mirus Bio #2700) and 100 ml of Opti-MEM per well. Lentivirus harvested at 72 hours was concentrated to 1/10^th^ of the total volume using the Lenti-HEK concentrator (Takara bio #631231). Virus was added to the HEK293T cell-line. Cells were plated at 50,000 in a 24-well plate and 30 µl of virus was added. Further, cells were moved up to a six-well plate after 48 hours and harvested, and PFA fixed for flow cytometry at 3, 5, 7 and 9 days. Partial UV photoconversion of the mEOS3.2 fluorophore was used to assess the amount of spontaneous aggregation by FRET using flowcytometry for each of the tau constructs. Fixed cells underwent partial UV photoconversion of the mEOS3.2 fluorophore for 30 minutes at 1.5 inches above light source (Chauvet LED Shadow - Model: TFX-UVLED) prior to flow cytometry.

For the seeding assays with tauRD mEOS3.2 constructs, 50,000 cells were plated in a 24-well plate and transduced after 18 hours with 30µl of virus. Further, cells were trypsinized after 72 hours and 10,000 cells were replated as triplicates in 96-well plates for exogenous seeding. Transduced cells were seeded exogenously with brain lysates from various tauopathies, lysates from PS19 tauopathy mice, and 2N4R tau recombinant fibrils (heparin induced) using lipofectamine 2000. After 72 hours, cells were harvested by 0.25% trypsin and then fixed in 1x PBS with 2% paraformaldehyde (Electron Microscopy Sciences #15710). Fixed cells were resuspended in 1x PBS and were run on flow cytometer to quantify the percentage of FRET positivity. The spontaneous aggregation of these 8 tauRD constructs was reproduced as two independent biological replicates (R1 and R2), each with three technical replicates. Direct comparison of the Pearson correlation for the percentage of FRET positive cells at 15,000 mEOS3.2_red_ intensity between values between R1 and R2 across all tauRD constructs at each time point (Days 3, 5, 7 and 9) to be 0.85-1.0 (Supplementary Fig. 9d, e).

### Preparation of brain tissue lysates for exogenous seeding in tauRD mEOS3.2 constructs

Frontal lobe sections of 0.5 g from AD, CBD, CTE and PSP patients, were gently homogenized at 4°C in 5 mL of TBS buffer containing cOmplete mini EDTA-free protease inhibitor cocktail tablet (Roche) at a concentration of 20% w/v using a Dounce homogenizer. Samples were centrifuged at 21,000 × *g* for 15 min at 4°C to remove cellular debris. Supernatant was partitioned into aliquots, snap-frozen in liquid nitrogen, and stored at -80°C. A similar process was used to generate PS19 mice brain lysates.

### Flow cytometry analysis

#### Peptide and tauRD seeding assay

The Attune CytPix/CytKick Flow cytometer from ThermoFisher Scientific was used to perform flow cytometry. For the peptide and tauRD seeding assay, mClover3, mCerulean3, and FRET were measured with the AmCyan channel. To measure mCerulean and FRET signal, cells were excited with the 405 nm laser, and fluorescence was captured with a 450/50mClover, cells were excited with a 488 laser, and fluorescence was captured with a 525/50 nm filter. To quantify FRET, we used a gating strategy where mCerulean3 bleed-through into the mClover3 and FRET channels was compensated using the Attune Software. The FRET gate was adjusted for biosensor cells that received lipofectamine alone and are thus FRET-negative. This allows for direct visualization of sensitized acceptor emission arising from the excitation of the donor at 405 nm. The FRET signal is defined as the percentage of FRET-positive cells in all analyses. For each experiment, 10,000 events per replicate were analyzed, and each condition was analyzed in triplicates. Experiments were carried out on Attune Cytpix flowcytometer (Thermo). Data analysis was performed using FlowJo v10 software (Treestar). Gating strategy for the analysis is in Supp Fig 4. Parameters used for flow cytometry are mentioned in Supplementary Table 2.

#### Spontaneous aggregation assay and tauRD seeding assay for tau RD mEOS3.2 constructs

The Attune CytPix/CytKick Flow cytometer from ThermoFisher Scientific was used to perform flow cytometry. For the mEOS tauRD spontaneous aggregation experiment, FRET between mClover3, and mRuby3 which is PerCP was measured. mEOS3.2 is a photoconvertible fluorescent protein which on partial photconversion forms a red-green FRET pair. Gating strategy for the spontaneous aggreagtion and seeding assay of tauRD mEOS3.2 constructs for analysis is shown in Supp Fig 9. Parameters used for flow cytometry are listed in Supplementary Table 3. Compensation was manually applied to correct for donor bleed-through into the FRET channel guided by a sample with non-aggregated and photoconverted tauRD-mEOS. Samples were gated on the acceptor intensity (mEOS3.2_red_ intensity) such that cells with similar concentrations of tauRD-mEOS were analyzed to mitigate the contribution of differences in concentration leading to apparent changes in the fraction of FRET-positive cells in each condition. mEOS3.2_red_ intensity binned as (5, 10, 15, 20, 25) k arbitrary units. For each experiment, 20,000 events per replicate were analyzed, and each condition was analyzed in triplicates. This experiment was reproduced with two independent biological replicates, each with three technical replicates. Comparison of percentage of cells that were FRET positive for each of the 7 tauRD constructs (calculated from population of cells with 15k mEOS3.2_red_ intensity populations) showed similar kinetics and a high Pearson correlation for between replicate 1 and replicate 2 (R1 and R2; Supplementary Fig. 9d,e). Data analysis was performed using FlowJo v10 software (Treestar). For tauRD mEOS3.2 constructs gating strtegy was similar to spontaneous aggregation assay.

#### Cell-based alanine scan for tauRD mEOS3.2 constructs

Cell based alanine scans were performed by plating 15,000 HEK293T cells in 96-well plates and concurrently treating with 10-20 uL lentivirus containing each tauRD alanine mutant with a target of >90% transduction for all mutants. Each well in the 96-well dish contained an individual tauRD alanine mutant. After 48 hours, the cells were replated into six new 96-well plates at a 1:6 ratio. AD, CBD, and PSP brain homogenates were optimized to maximize percentage of cells with aggregates in WT tauRD cell lines while minimizing cell toxicity. This resulted in treatments ranging from 10 to 25 μg of 10% w/v brain homogenate. Samples were prepared in a similar manner as described above with 0.75 ul/well of Lipofectamine 2000 and variable Opti-MEM to maintain a total treatment volume of 50uL per well. After incubation for 48 to 72 hours, the cells were harvested by 0.25% trypsin digestion for 5 min at 37C, quenched and resuspended with cell media, transferred to 96-well U-bottom plates, and centrifuged for 5 min at 200g. The cells were then fixed in 1x PBS with 2% paraformaldehyde for 10 min before a final centrifugation step and resuspension in 150 μl of 1x PBS. Fixed cells underwent partial UV photoconversion of the mEOS3.2 fluorophore for 30 minutes at 1.5 inches above light source (Chauvet LED Shadow - Model: TFX-UVLED) prior to flowcytometry analysis using BD-LSR Fortessa SORP instrument. To characterize the fluorescence of mEOS3.2, cells were analyzed in both green and red states using specific laser-filter combinations. Green-state emission was captured via 488-nm excitation paired with a 525/50-nm band-pass filter, while the photoconverted red state was detected using 561-nm excitation and a 610/20-nm filter. FRET was quantified by exciting the donor at 488 nm and monitoring emission through the 610/20-nm filter which is denoted as the PerCP channel. Single HEK293 cells were identified using FSC/SSC profiles. Aggregate-positive cells were isolated using FRET in the PerCP channel. Analytical gates were established based on a suite of negative and positive controls, including wild-type tauRD, tauRD I277P_I308P (negative for aggregates), mEOS3.2 monomer, and the mEOS3.2-linker-mEOS3.2 construct. To ensure the acquisition of consistent raw values for downstream statistical comparison, all gating boundaries and thresholds were maintained identically across all tau variants and experimental replicates.

### Immunoblotting for tauRD mEOS3.2 constructs

Transduced cells were harvested on Day 9, endpoint of our spontaneous aggregation assay for our tauRD S320F and tauRD S320F cysteine mutants. Harvested cell pellets were washed twice with ice-cold 1x PBS and further frozen in -20°C later usage. For immunoblotting with tau, cell lysates were prepared in 1% TritonX100 in 1X TBS containing cOmplete mini EDTA-free protease inhibitor cocktail tablet (Roche). Cells were subjected to mechanical shear using a 26-gauge syringe followed by pre-clearance at 500 × *g* for 30 min at 4°C. Further, the supernatant from previous step was used to do fractionation at 186,000 × *g* using ultracentrifugation. Later, pellets from ultracentrifugation were resuspended in 200 µl of 1% Triton-X-100 in 1X TBS. Both resuspended pellet and supernatant were quantified for total protein abundance using BCA (Bicinchoninic Acid) Assay (Pierce#A55865). Samples were prepared in LDS sample buffer with and without β-ME and boiled for 5 min at 95°C prior to loading on gel. 6 µg of total protein was loaded on pre-casted 4-12% NuPAGE Bis-Tris gels (Invitrogen#WG1403). Gels were run at 150V for 1hour 30 min. For western blotting, semi-dry transfer was performed using 0.2 µm PVDF membrane for 1 hour at 20V using Novoblot transfer equipment. Membranes were blocked using 5% blotting-grade milk in 1X TBS buffer with 0.01% Tween-20 for 2 hours. Membranes were incubated with a tau-A antibody that recognizes epitopes 246-266 in the tau repeat domain (from Marc Diamond’s Lab at UTSW) at the dilution of 1:10,000 at 4°C with overnight shaking, followed by incubation with anti-mouse secondary-HRP conjugated antibody. Blots were developed using G-Biosciences femtoLUCENT^TM^ PLUS.

### Fluorescence microscopy for tauRD mEOS3.2 constructs

For imaging, cells were plated at a confluency of 50,000 cells in each well of cell-culture treated 24-well plates. After 18 hours, cells were transduced with 50 ml of tauRD mEos3.2 constructs virus. Cells were transduced for 48 hours and then harvested, and 15,000 cells were replated in a fibronectin-coated 24-well coated glass-bottom plate (Cellvis #P24-1.5H-N). Prior to imaging Hoechst stain (Invitrogen# 33342, stock of 10 mg/ml) was added to the wells at the concentration of 1 mg/ml incubated at RT for 15 min. Live-cell imaging was performed using a Nikon SORA (David Sander’s Lab, UTSW) microscope. TauRD S320F, and tauRD S320F C291S were imaged at day 3. For the rest of the tauRD mEOS3.2 constructs, imaging was performed at day 9 after transduction. DAPI and GFP channel images were acquired at 60X magnification with an oil-immersion lens. Images were exported in nd2 format for downstream analysis. Single color panels were merged, and composite images were created using Fiji software^72^.

## Data availability

The structural datasets generated during the current study are available in the Electron Microscopy Data Bank repository (https://www.ebi.ac.uk/emdb/) under accession number, EMD-71887 [https://www.ebi.ac.uk/emdb/EMD-71887]. The structural model that was fit into the density is available in the Protein Data Bank under PDB id 9PVA (https://www.rcsb.org/structure/unreleased/9PVA). ThT aggregation data, ddG calculations and cell-based aggregation data are available as Source Data. The raw data associated with these data are available on Zenodo under accession number 18665770. Any other data sets generated during and/or analyzed during the current study are available from the corresponding author on reasonable request. PBD id’s used in this study are: 6hre [http://doi.org/10.2210/pdb6HRE/pdb], 6nwp [http://doi.org/10.2210/pdb6NWP/pdb], 6tjo [http://doi.org/10.2210/pdb6TJO/pdb] and 7p65 [http://doi.org/10.2210/pdb7P65/pdb].

## Code Availability

The software used to analyze and determine a cryo-EM structure of the S320F_295-330_ fibril included CTFFIND v4.1^66^, RELION v4.0^45^ [https://www3.mrc-lmb.cam.ac.uk/relion/index.php/Main_Page], COOT v9.4^46^ [https://www2.mrc-lmb.cam.ac.uk/personal/pemsley/coot/]. All ΔΔG calculations on the cryo-EM fibril structure was performed using an adapted flex_ddg protocol^22^ available on [https://git.biohpc.swmed.edu/s184069/flex_ddg_ala_scn_runner] using Rosetta v3.12 [https://www.rosettacommons.org/]. Flow cytometry data was analyzed using FlowJo v10 [https://www.flowjo.com/solutions/flowjo].

## Acknowledgements

The U.S. Army Medical Research Acquisition Activity, 808 Schreider Street, Fort Detrick MD 21702-5014 is the awarding and administering acquisition office and this work was supported by The Assistant Secretary of Defense for Health Affairs endorsed by the Department of Defense, in the amount of $1,639,837 through the Peer Reviewed Medical Research Program under Award Number HT9425-24-1-0641. Opinions, interpretations, conclusions, and recommendations are those of the author(s) and are not necessarily endorsed by The Assistant Secretary of Defense for Health Affairs endorsed by the Department of Defense. In the conduct of research utilizing recombinant DNA, the investigator(s) adhered to NIH Guidelines for research involving recombinant DNA molecules. L.A.J. was additionally supported by an Effie Marie Cain Scholarship in Medical Research and a grant from the NIH-NIA (RF1AG076459). M.I.D. was supported by a grant from NIH-NIA (RF1AG065407). Transmission electron microscopy was performed at the UT Southwestern Electron Microscopy Core Facility, supported by the National Institutes of Health (1S10OD021685-01A1 and 1S10OD020103-01). Mass Spectrometry was performed at the UT Southwestern Proteomics Core Facility. Cryo-EM data were acquired at the Cryo-Electron Microscopy Facility (CEMF) and the Structural Biology Laboratory (SBL) at UT Southwestern, which are supported by a grant from the Cancer Prevention & Research Institute of Texas (RP220582). Computational resources were provided by the BioHPC cluster supported by the Lyda Hill Department of Bioinformatics at UT Southwestern. We thank Drs. Sanders and Banerjee for guidance on cell imaging using the Nikon SoRa. We thank members of the Joachimiak laboratory for helpful discussions and feedback on the manuscript.

## Author Contributions

P. J and L.A.J. initiated the project. Peptide and tau repeat domain aggregation experiments and their validation using TEM were performed by P. J. Cryo-EM structure of the S320F_295-330_ tau fibril was determined by P.J. and B.A.N. Spontaneous aggregation and seeding experiments were performed by S. R. using tauopathy tissues from C.L.W. Seeding with tau alanine mutants in cells was performed by J.V.A. and V.B., using tauopathy patient samples collected and characterized by C.L.W. L. S., S. S. and M.I.D. provided guidance on structure and cell experiments. Finally, L.A.J. conceived of and directed the research as well as wrote the manuscript. All authors contributed to the revisions of the manuscript.

---

**Supplementary Table 1.**
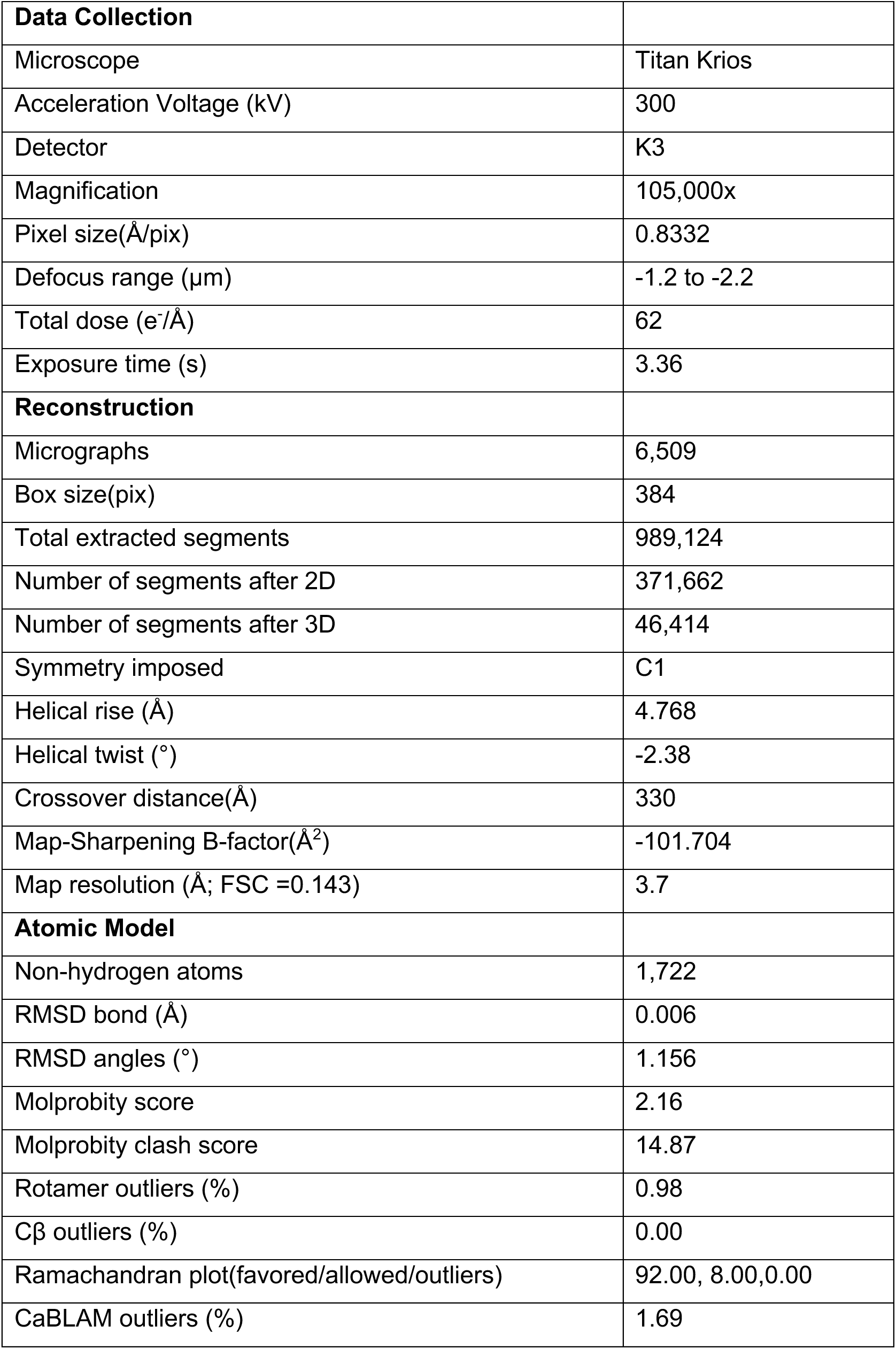
Cryo-EM data collection parameters for S320F_295-330_ tau fragment fibrils.

|  |  |
| --- | --- |
| **Data Collection** |  |
| Microscope | Titan Krios |
| Acceleration Voltage (kV) | 300 |
| Detector | K3 |
| Magnification | 105,000x |
| Pixel size(Å/pix) | 0.8332 |
| Defocus range (µm) | -1.2 to -2.2 |
| Total dose (e<sup>-</sup>/Å) | 62 |
| Exposure time (s) | 3.36 |
| **Reconstruction** |  |
| Micrographs | 6,509 |
| Box size(pix) | 384 |
| Total extracted segments | 989,124 |
| Number of segments after 2D | 371,662 |
| Number of segments after 3D | 46,414 |
| Symmetry imposed | C1 |
| Helical rise (Å) | 4.768 |
| Helical twist (°) | -2.38 |
| Crossover distance(Å) | 330 |
| Map-Sharpness B-factor(Å<sup>2</sup>) | -101.704 |
| Map resolution (Å; FSC =0.143) | 3.7 |
| **Atomic Model** |  |
| Non-hydrogen atoms | 1,722 |
| RMSD bond (Å) | 0.006 |
| RMSD angles (°) | 1.156 |
| Molprobit score | 2.16 |
| Molprobit clash score | 14.87 |
| Rotamer outliers (%) | 0.98 |
| Cβ outliers (%) | 0.00 |
| Ramachandran plot(favored/allowed/outliers) | 92.00, 8.00,0.00 |
| CaBLAM outliers (%) | 1.69 |

**Supplementary Table 2:** Parameters used during flow cytometry data collection for peptide and tauRD seeding assays in NK13 (tauRD P301S biosensors)

| Collected parameter | Voltage | Excitation (nm) | Emission filter (nm) |
| --- | --- | --- | --- |
| Forward Scatter | 100 | 488 |  |
| Side Scatter | 380 | 488 |  |
| Acceptor, "mClover3" | 220 | 488 | 525/50 |
| Donor, "mCerulean3" | 300 | 405 | 450/50 |
| FRET, "AmCyan" | 200 | 405 | 525/50 |

**Supplementary Table 3:** Parameters used during flow cytometry data collection of tauRD-mEOS3.2 expression system.

| **Collected parameter** | Voltage | Excitation (nm) | Emission filter |
| --- | --- | --- | --- |
| Forward Scatter | 100 | 488 | - |
| Side Scatter | 380 | 488 | 488/10 BP |
| Acceptor, “Alexa Fluor 488” | 220 | 488 | 530/30 |
| Donor, “mRuby3” | 300 | 561 | 620/15 |
| FRET, “PerCP” | 300 | 488 | 695/40 |

**Supplementary Table 4:** Subjects studied.

| Sample Name | ID | Age | Sex | Institution |
| --- | --- | --- | --- | --- |
| AD_46121 | 46121 | 74 | F | UTSW |
| AD_46879 | 46879 | 76 | M | UTSW |
| AD_49341 | 49341 | 83 | F | UTSW |
| AD_47833 | 47833 | 82 | M | UTSW |
| AD_46090 | 46090 | 75 | F | UTSW |
| AD_45408 | 45408 | 73 | M | UTSW |
| AD_62579_Parietal | 62579 | 71 | M | WashU |
| AD_63007_Parietal | 63007 |  |  | WashU |
| AD_63870_Parietal | 63870 | 81 | F | WashU |
| AD_63870_Temporal | 63870 | 81 | F | WashU |
| CBD_48638 | 48638 | 64 | M | UTSW |
| CBD_44193 | 44193 | 55 | M | UTSW |
| CBD_46267 | 46267 | 67 | F | UTSW |
| CBD_3950 | 3950 | 72 | M | WashU |
| CBD_23302 | 22302 | 62 | M | WashU |
| CTE_8240 | 9130 | 40 | M | Boston University |
| CTE_6227 | 8408 | 49 | M | Boston University |
| CTE_9284 |  |  |  | Boston University |
| CTE_7865 |  |  |  | Boston University |
| PSP_45460 | 45460 | 68 | F | UTSW |
| PSP_47941 | 47941 | 69 | M | UTSW |
| PSP_49017 | 49017 | 61 | F | UTSW |
| PSP_48221 | 48221 | 67 | M | UTSW |

**Supplementary Figure 1.**
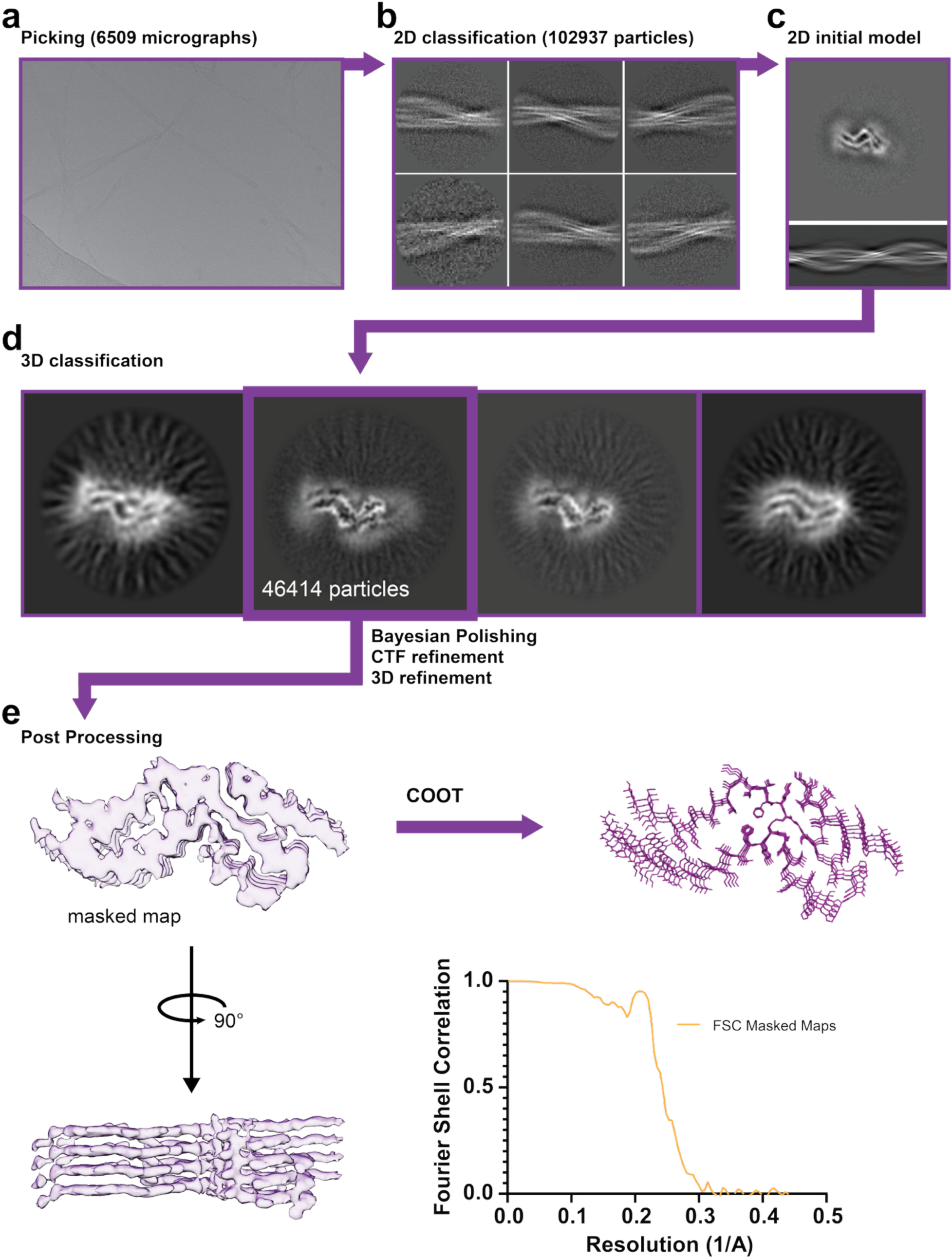
Cryo-EM reconstruction of S320F_295-330_ tau fibrils. Representative micrograph (**a**) and 2D class averages (**b**) were used to generate an initial 2D reference model (**c**). After particle selection (46,414 particles), multiple 3D classifications were performed (**d**), yielding a well-defined fibril reconstruction. The final 3D density map (**e**) was refined to high resolution, allowing atomic model building (top right) and visualization of fibril packing (bottom left inset). Fourier shell correlation (FSC) curves (bottom right) demonstrate the resolution and quality of the final reconstruction.

**Supplementary Figure 2.**
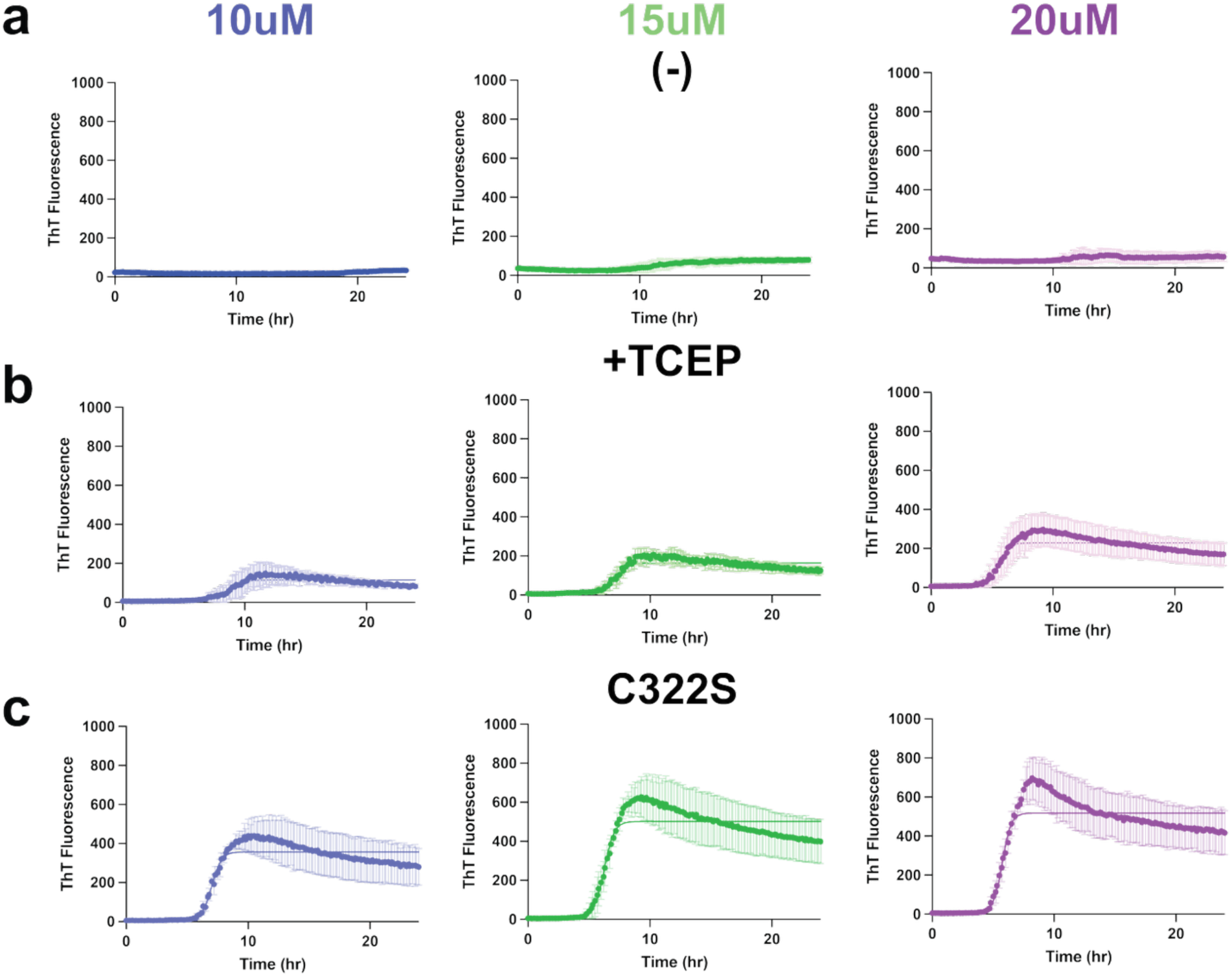
Kinetic traces of S320F_295-330_ and mutant peptide aggregation. ThT fluorescence aggregation kinetics on **(a)** S320F_295-330_ (-), **(b)** bS320F_295-330_ (TCEP), and **(c)** S320F_295-330_ C322S at three different concentrations; 10 µM (blue), 15 µM (green) and 20 µM (magenta). All experiments were carried out as technical triplicates and each curve was fit to a non-linear regression model in GraphPad Prism to estimate t_1/2_max and fluorescence amplitudes. The data is shown as an average with a standard deviation.

**Supplementary Figure 3.**
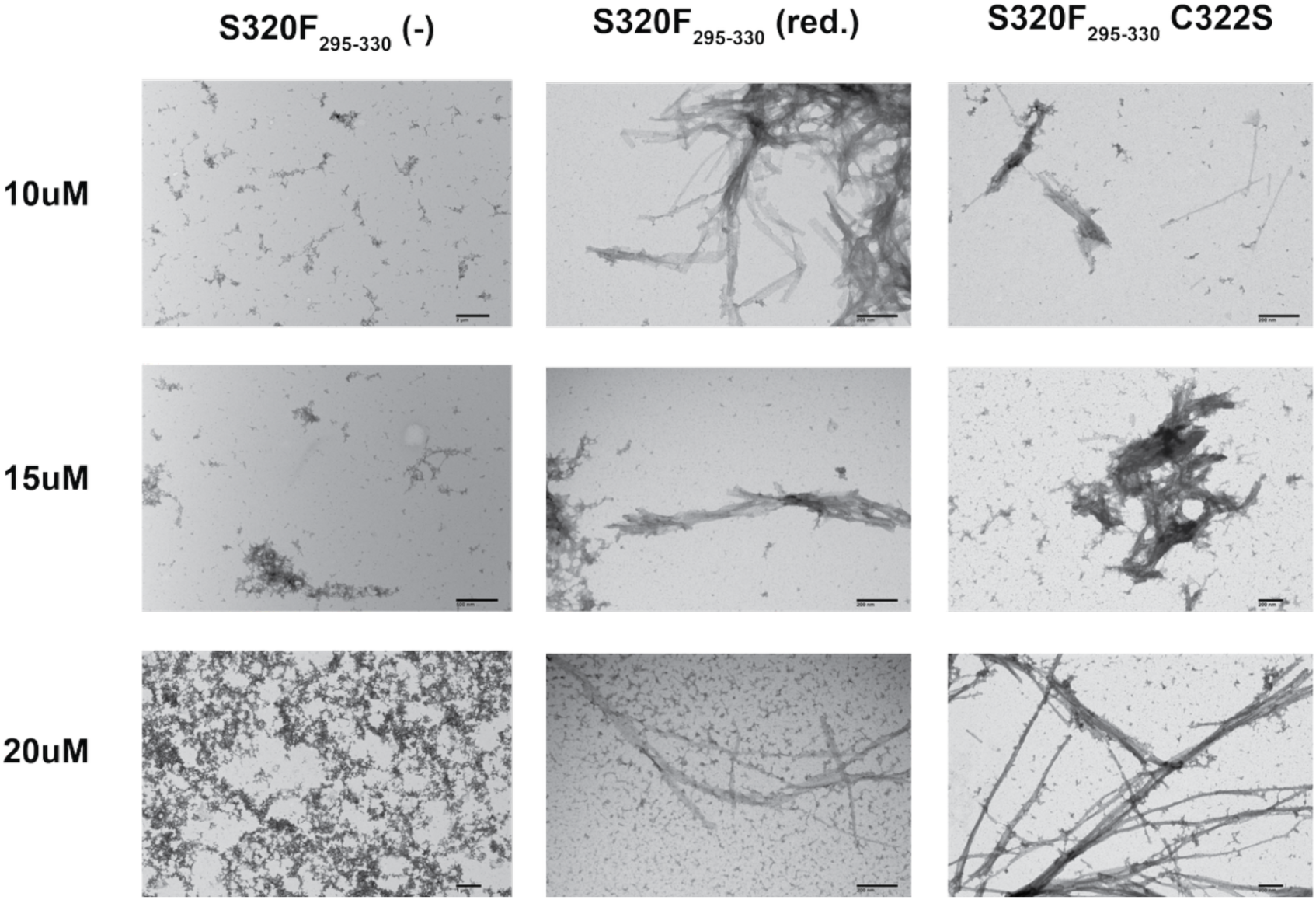
TEM verification of S320F_295–330_ and C322 variant fibril samples. TEM images at the endpoint of ThT fluorescence assay of S320F_295-330_ without TCEP (-), with TCEP (red.) and S320F_295-330_ C322S; 10 µM,15 µM and 20 µM (left to right). Scale bar represents 200 nm.

**Supplementary Figure 4.**
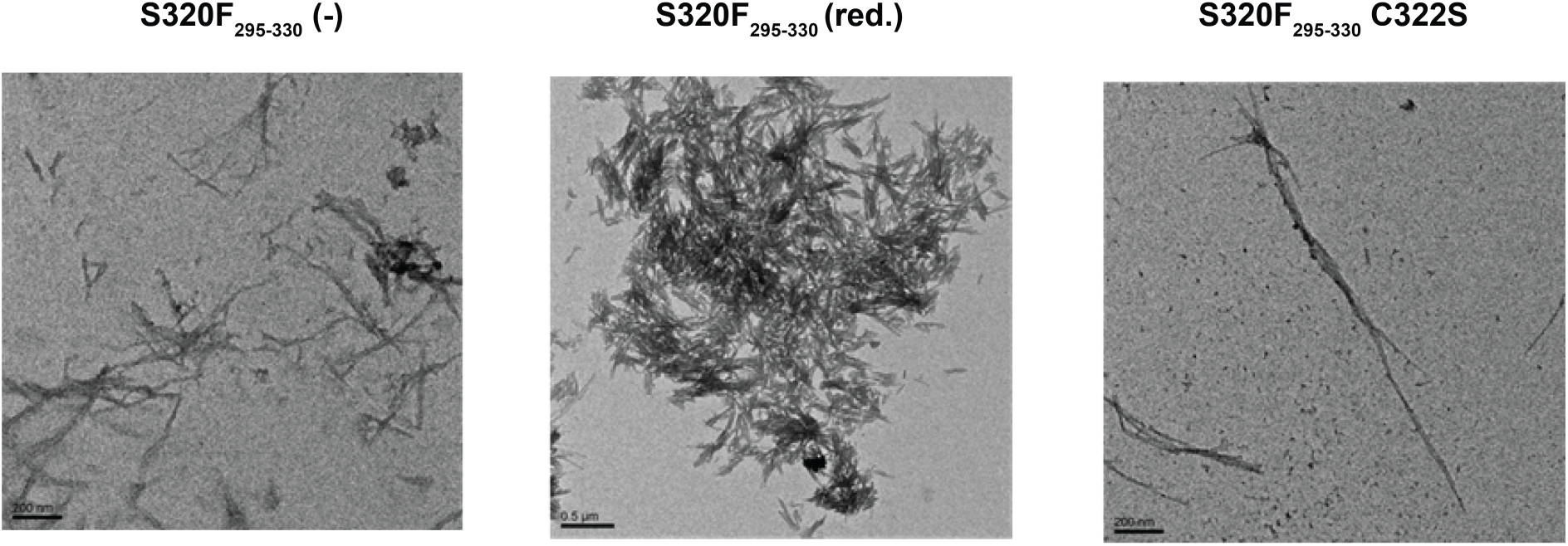
Biochemical characterization and aggregation properties of S320F tau peptides and C322 variant. Negative-stain TEM images of fibrils formed by S320F_295–330_ without TCEP (-), S320F_295–330_ with TCEP (red.), and cysteine variant (C322S) of S320F_295–330_. Scale bars 200-500 nm.

**Supplementary Figure 5.**
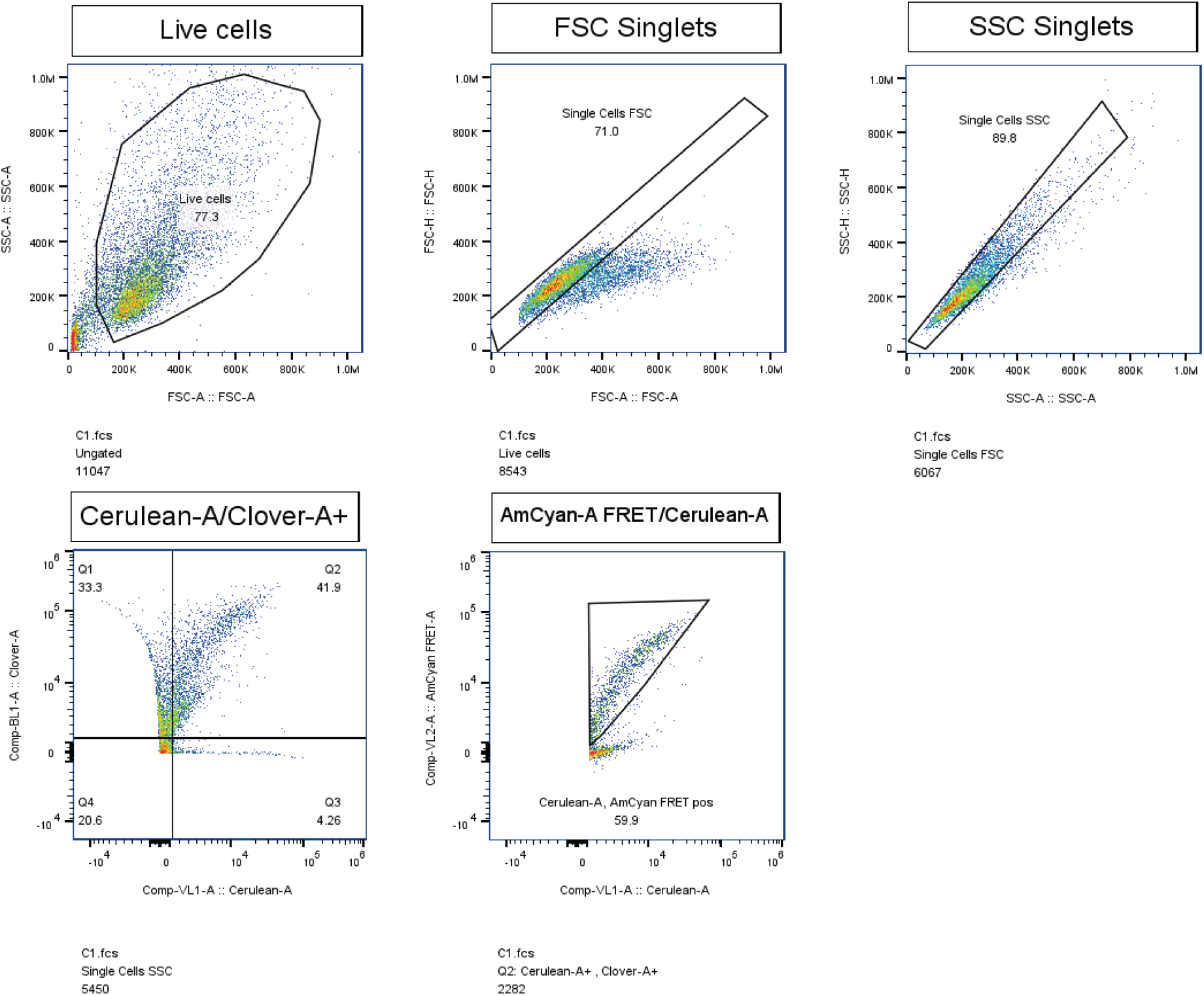
Gating strategy for seeding assay in tauRD P301S Cerulean/Clover biosensors with S320F peptides in various conditions and S320F tauRD, and S320F tauRD cysteine mutant fibrils. Gating strategy to extract live, single, mCerulean3 (cyan) and mClover3 (green) channel double-positive cells and expression level for FRET quantification between AmCyan-A FRET/mCerulean3-A.

**Supplementary Figure 6.**
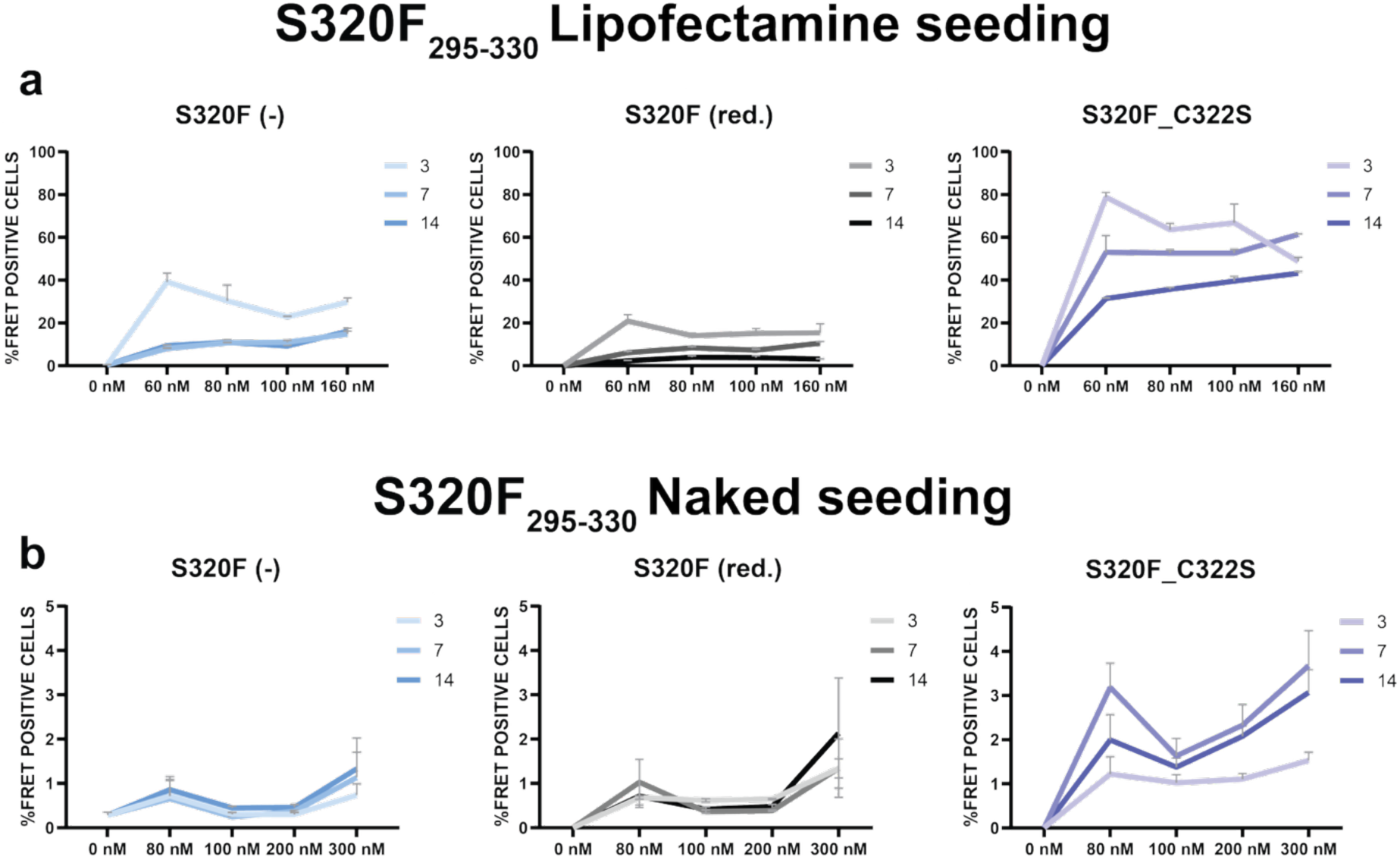
Titration of S320F_295-330_ and variant aggregates in cell seeding assays. (**a**) Lipofectamine-based seeding of fibrils S320F_295-330_ under non-reducing conditions (-) (no TCEP), S320F_295-330_ under reducing conditions (red.) (with TCEP) and S320F_295-330_ C322S. Seeding was performed across samples pulled at 3, 7 and 14 days at 5 concentrations of fibrils (0, 80, 100, 200, and 300) nM for naked seeding, (0, 60, 80, 100, and 160) for lipofectamine seeding. (**b**) Naked-based seeding (direct) of fibrils (d) S320F_295-330_ under non-reducing conditions (-) (no TCEP), S320F_295-330_ under reducing conditions (red.) (with TCEP) and S320F_295-330_ C322S. Seeding was performed across samples pulled at 3, 7 and 14 days at 5 concentrations of fibrils (0, 80, 100, 200, and 300 nM). Plots for the 3-day timepoint for S320F_295-330_ (-), S320F_295-330_ (red.), and S320F_295-330_ C322S samples are colored light blue, grey and violet, respectively with the 7-and 14-day time points colored as darker shades of blue, grey and violet. Data is shown as averages of triplicates with SEM.

**Supplementary Figure 7.**
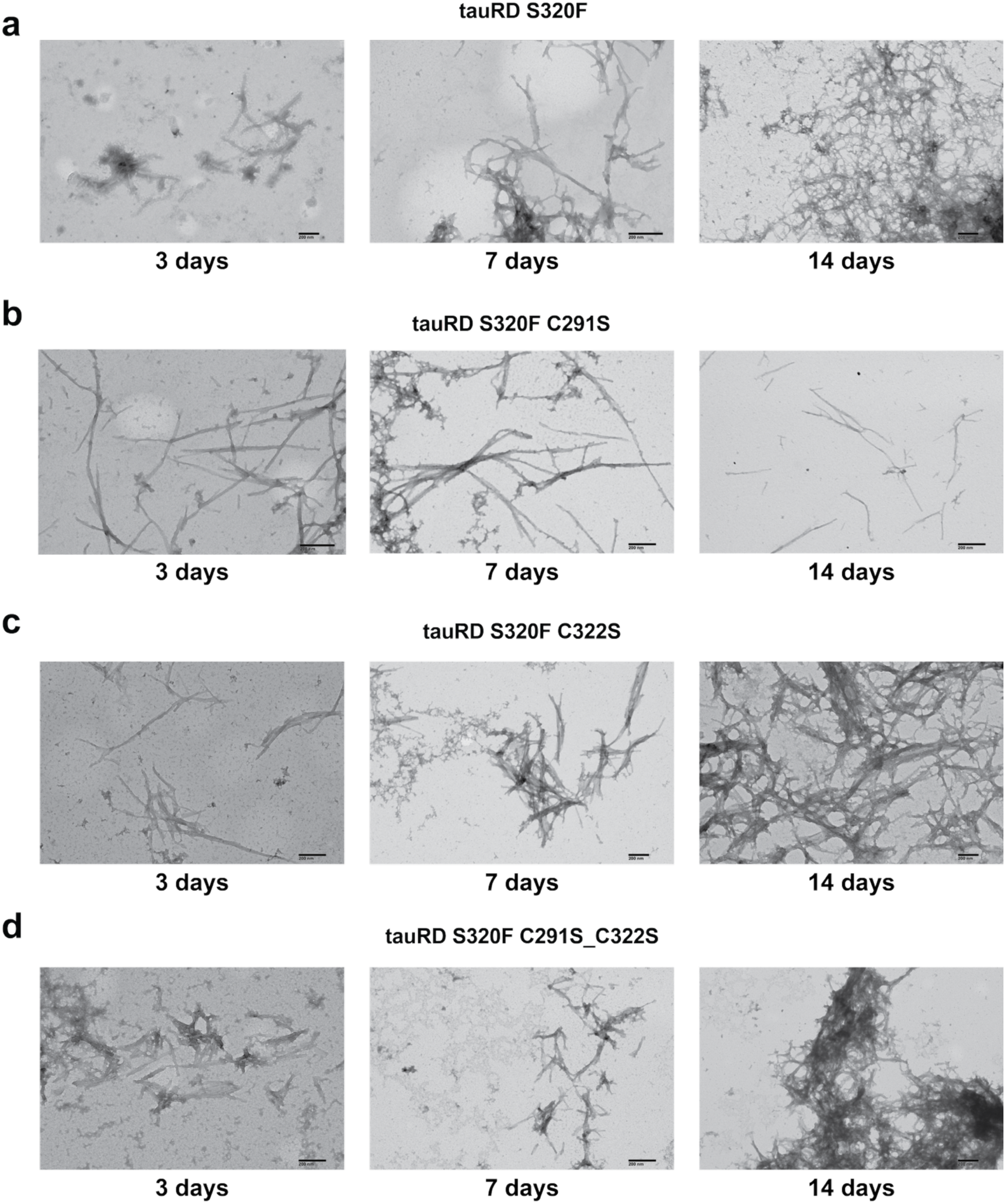
TEM-based confirmation of tauRD S320F, tauRD S320F C291S, tauRD S320F C322S and tauRD S320F C291S C322S aggregates. TEM images of fibrils samples used for the lipofectamine and naked seeding experiments with three different time points; 3, 7 and 14 days (left to right). (**a**) tauRD S320F (TCEP), (b) tauRD S320F C291S (TCEP), (**c**) tauRD S320F C322S (TCEP) and (**d**) tauRD S320F C291S_C322S (TCEP). Scale bar: 200 nm.

**Supplementary Figure 8.**
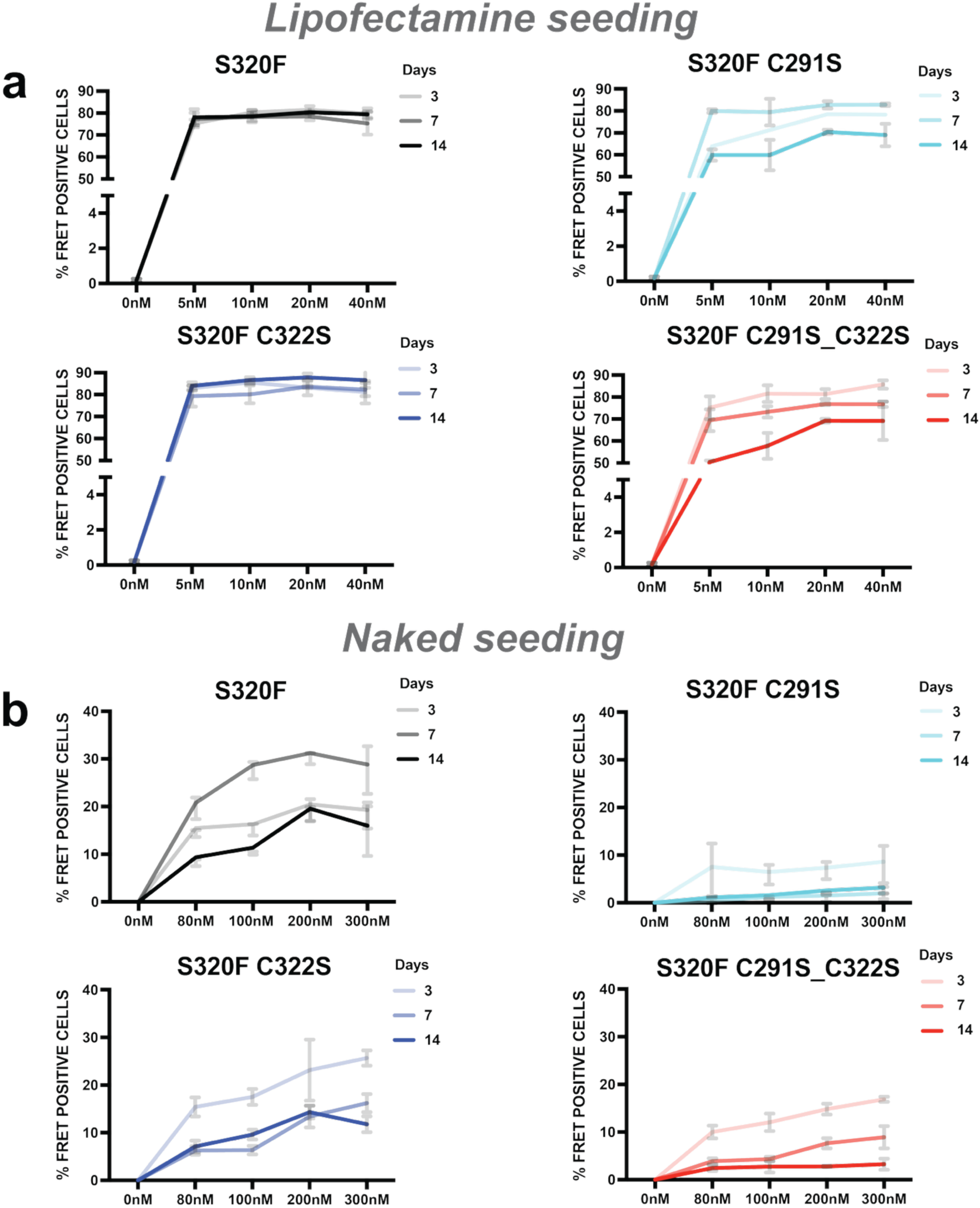
Cell-based seeding activity of *in vitro* tauRD S320F and tauRD S320F cysteine mutant fibrils. Seeding was evaluated for samples incubated in vitro for 3, 7 and 14 days and delivered into tau biosensor cells with (lipofectamine) or without transfection reagent (naked seeding). tauRD S320F in vitro purified constructs consists of amino acid residues from 243 to 378 with a mutation at 320 positions with two cysteines at positions C291 and C322. Seeding experiments were carried out with fibrils at three time points (3, 7 and 14 days) using naked and lipofectamine mediated seeding. (**a**) Percent FRET positivity from lipofectamine-mediated seeding activity of fibrils formed from tauRD S320F (TCEP) (black), S320F C291S (cyan), S320F C322S (blue) and S320F C291S_C322S (red). Preformed fibrils used are 0, 5, 10, 20, and 40 nM concentrations incubated with lipofectamine. 0nM is lipofectamine (vehicle only control). (**b**) Percent cell FRET positivity from naked seeding activity (in the absence of transfection reagents) of fibrils formed by incubating tauRD S320F (black), S320F C291S (cyan), S320F C322S (blue) and S320F C291S_C322S (red) preformed fibrils at 0, 80, 100, 200 and 300 nM concentrations with cells. All data is in triplicates tittered from 80nM to 300nM. All the values plotted are averages with SEM. Data is shown as averages of triplicates with SEM.

**Supplementary Figure 9.**
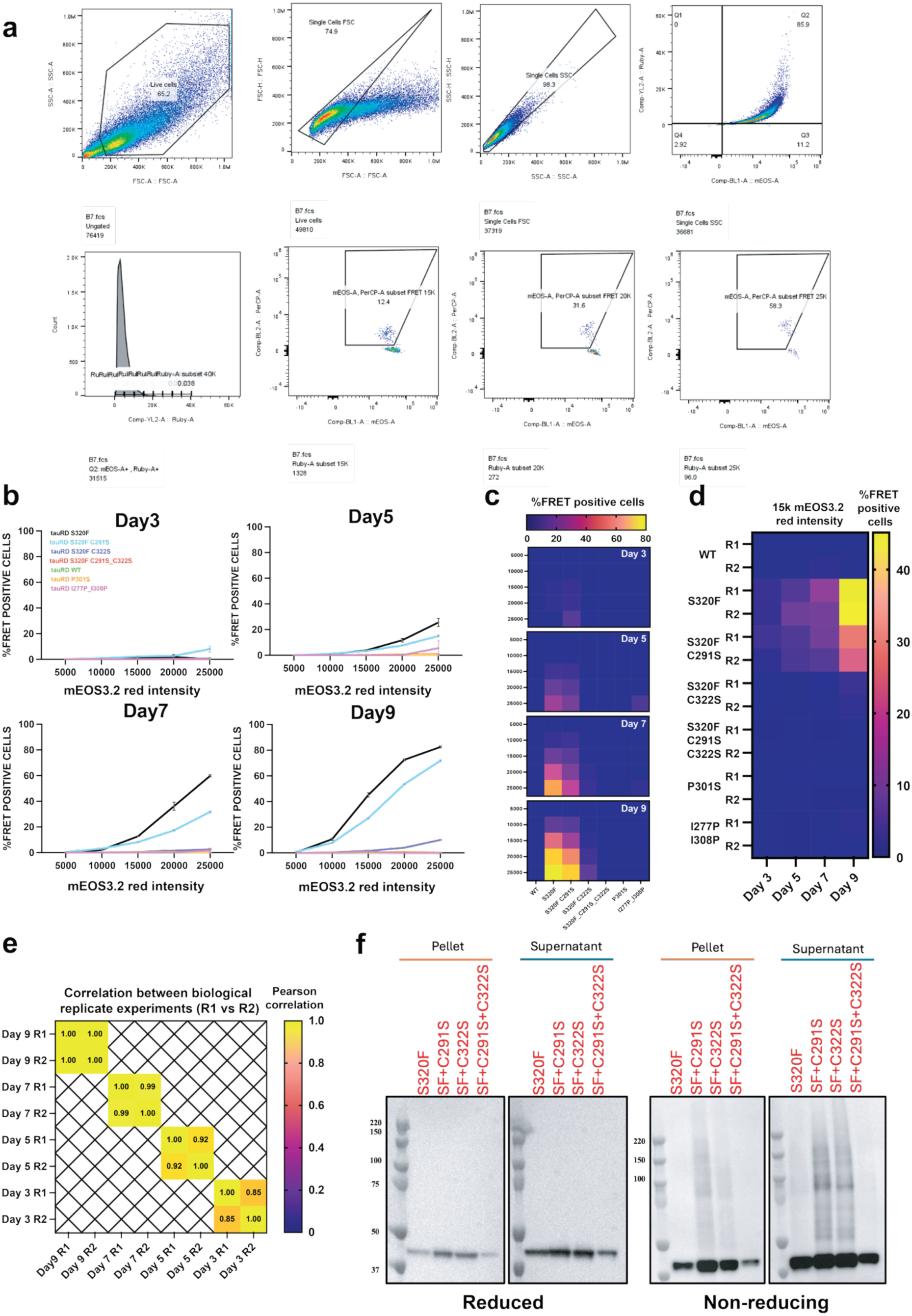
Time course to characterize S320F tauRD and cysteine mutant spontaneous aggregation in cells. **(a)** Representative flow cytometry gating strategy and distribution plots used to quantify aggregate-positive cells, showing sequential gating, fluorescence intensity thresholds, and population shifts upon spontaneous aggregation. **(b)** Time-course quantification of aggregate accumulation, plotted as mean signal intensity on x axis and percentage of FRET positive cells on y-axis across multiple days (Days 3, 5, 7, and 9). Data is shown as averages of three technical replicates with standard error of mean . **(c)** Heat maps summarizing spontaneous in cell aggregation kinetics across tauRD S320F and tauRD S320F cysteine mutants and time points, highlighting progressive increases in percentage of FRET positive cells as a function of red fluorescence signal intensity (5k, 10k, 15k, 20k, 25k) in arbitrary units. Data is shown as average from three technical replicates. Heatmaps are colored in plasma, with yellow High FRET and blue Low FRET. (**d**) Comparison of replicate 1 (R1) and replicate 2 (R2), biological replicates for spontaneous aggregation of tauRD S320F, tauRD S320F C291S, tauRD S320F C322S, tauRD S320F C291S_C322S, tauRD P301S, tauRD WT and tauRD I277P_I308P measured at days 3, 5, 7 and 9. Data is shown as a heat map comparing percentage of cells with FRET positive aggregates (from 15k mEOS3.2_red_ intensity) for each construct in R1 and R2. Data is shown as an average across three technical replicates. Heat map is colored in plasma with high and low percentage of cells with FRET positive cells colored yellow and blue. (**e**) Pearson correlation of percentage of FRET positive cells (at 15k mEOS3.2_red_ intensity) for replicate 1 (R1) and replicate (R2) across all constructs at days 3, 5, 7 and 9. Data is shown as a heat map and colored from yellow for high correlation and red for low correlation. **(f)** Biochemical validation of assemblies immunoblotting against tau of pellets and supernatants of lysates from day 9 after transduction resolved on SDS-PAGE under reducing and non-reducing conditions.

**Supplementary Figure 10.**
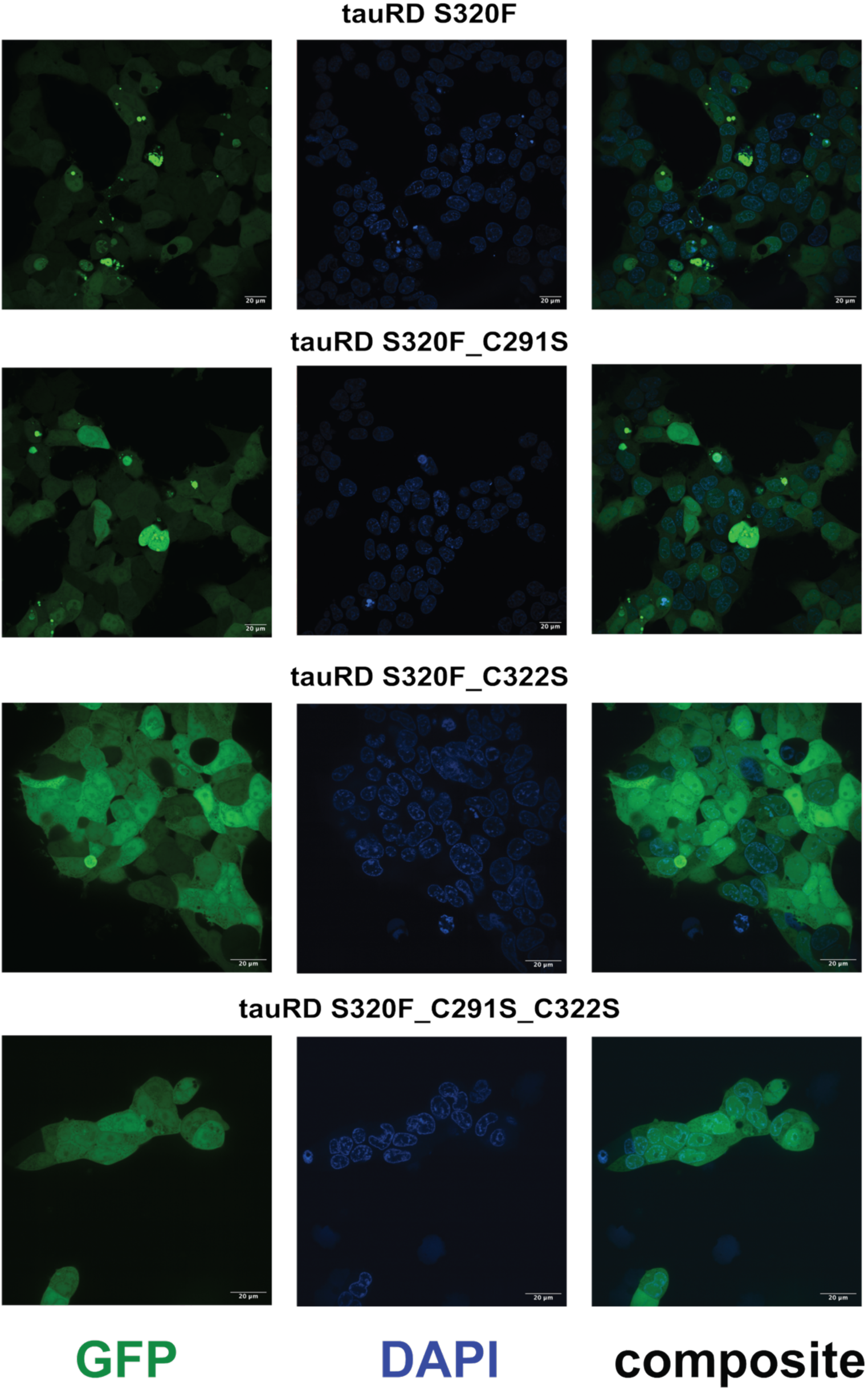
Effects of cysteine mutations on spontaneous aggregation of tauRD S320F in cells. Representative fluorescence microscopy images of cells expressing repeat domain (tauRD) S320F and cysteine variants. Rows show tauRD S320F (top), tauRD S320F C291S, tauRD S320F C322S, and tauRD S320F C291S_C322S (bottom). Columns display GFP fluorescence (left; tau aggregation reporter), DAPI nuclear staining (middle), and merged composites (right). Robust GFP-positive inclusions are observed for tauRD S320F and the C291S variant, indicating efficient aggregation and seeding in cells. By contrast, mutation of C322S blocks and double cysteine mutant (C291S_C322S) strongly suppress inclusion formation. Scale bars, 20 μm. Cells were imaged at 9 days post transduction, except for tauRD S320F and tauRD S320F C291S which were imaged at day 3. All images were processed using Fiji.

**Supplementary Figure 11.**
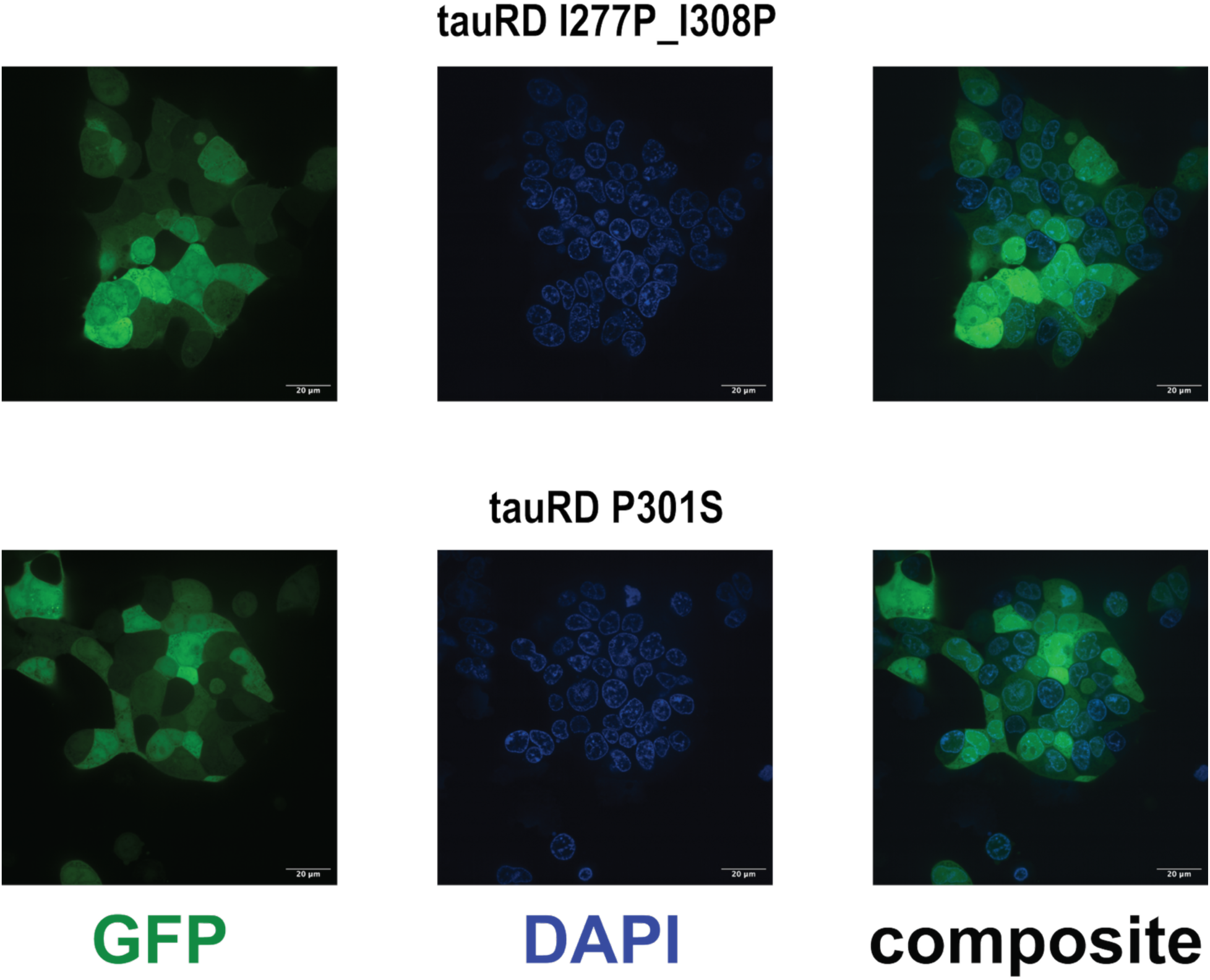
Cellular aggregation phenotypes of control I277P_I308P and P301S tauRD constructs. Representative fluorescence microscopy images of cells expressing tau repeat domain (tauRD) variants encoding either the double proline mutation I277P_I308P (top row) or the frontotemporal dementia-associated mutation P301S (bottom row). GFP fluorescence (left) reports tauRD, DAPI (middle) labels nuclei, and merged composite images (right) show the spatial relationship between diffusely localized tauRD and cell nuclei. Scale bars, 20 µm. Cells were imaged at 9 days post transduction. All images were processed using Fiji.

**Supplementary Figure 12.**
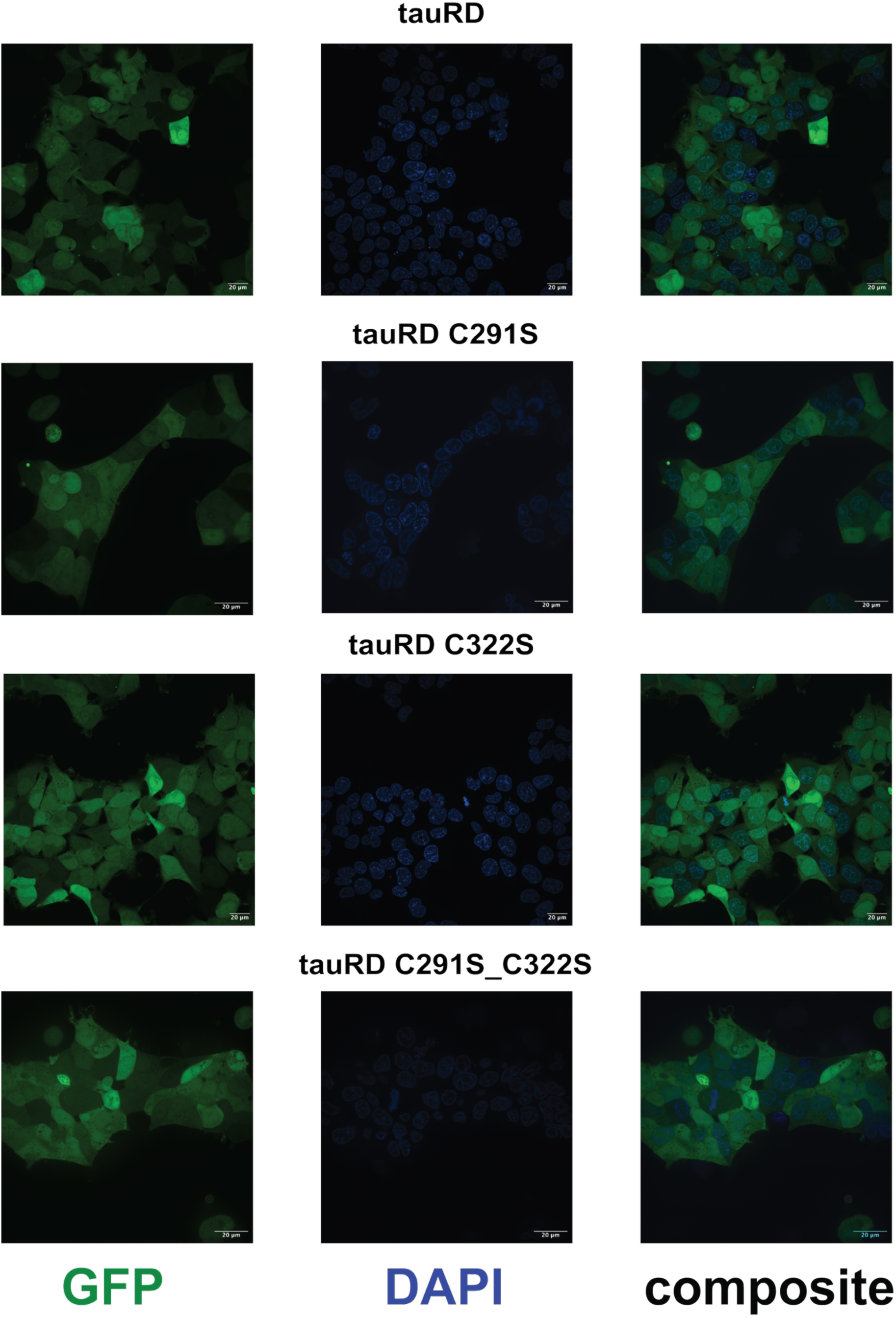
Cysteine-dependent modulation of tauRD inclusion formation. Representative fluorescence microscopy images of cells expressing the tau repeat domain (tauRD) and cysteine variants. Rows show tauRD (WT) (top), tauRD C291S, tauRD C322S, and the double mutant tauRD C291S_C322S (bottom). GFP fluorescence (left) reports tauRD inclusion formation, DAPI staining (middle) labels nuclei, and merged composite images (right) show the cellular distribution of tauRD relative to nuclei. WT tauRD, tauRD C291S, tauRD C322S and tauRD C291S_C322S do not form inclusions over the course of this experiment. These images demonstrate that native cysteine residues modulate tauRD self-association and aggregation behavior in a cellular context. Scale bars, 20 µm. Cells were imaged at 9 days post transduction. All images were processed using Fiji.

**Supplementary Figure 13.**
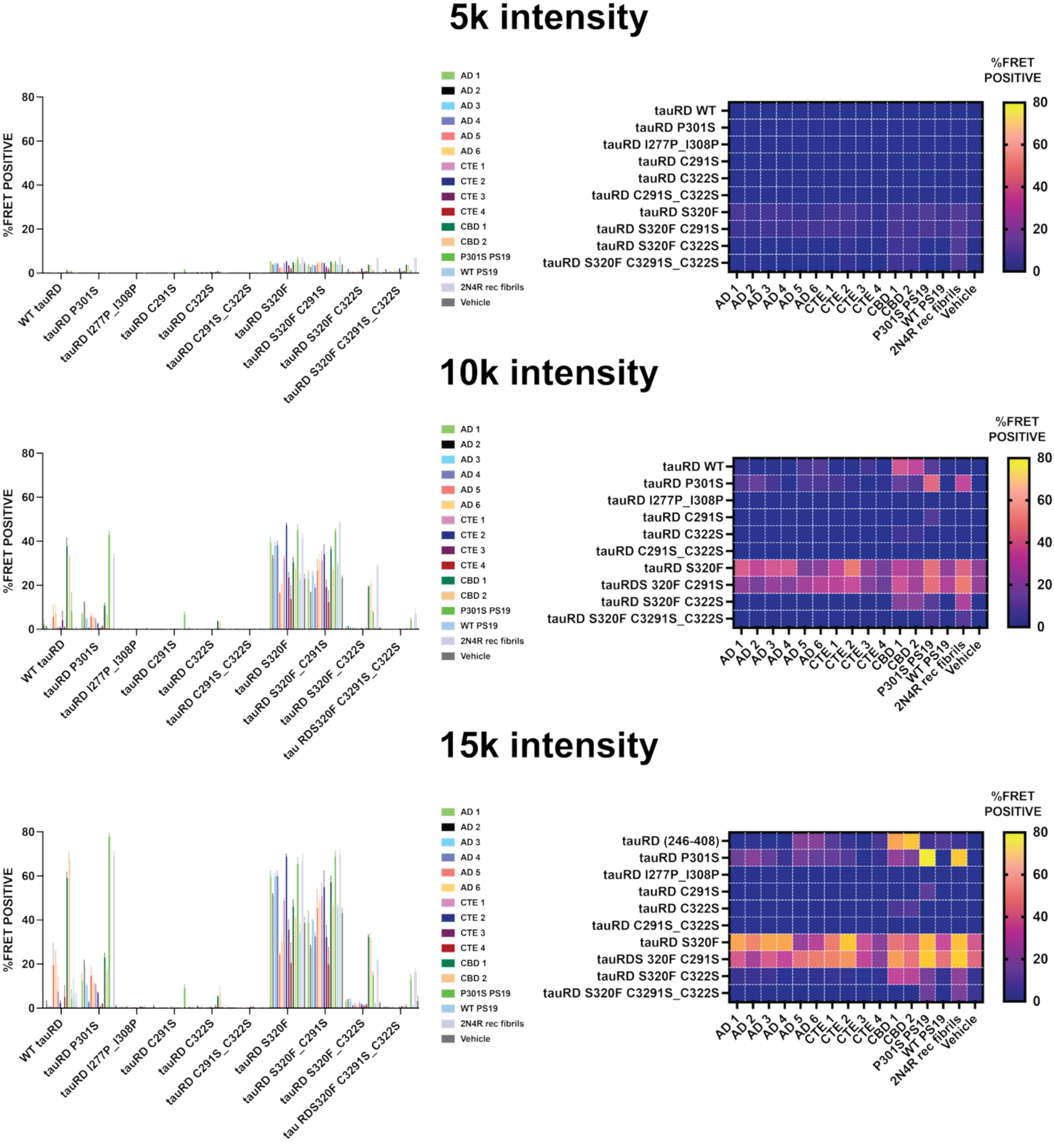
Seeded aggregation of cysteine mutants in WT tauRD and tauRD S320F. Analysis comparing WT tauRD and S320F tauRD constructs encoding cysteine mutations, illustrating differential seeded aggregation propensities. Spontaneous aggregation shown separately for WT tauRD or tauRD S320F with cysteine mutants at 5K (top), 10K (middle), 15K (bottom) at mEOS3.2_red_ intensity. Cells were seeded with tauopathy tissues including AD, CTE, CBD, PS19 and PS19 control animal tissues and recombinant 2N4R tau heparin aggregates. Experiments were performed in triplicate with data shown as average with standard error of mean in bar plot representation (left) and averages as heatmaps shown in plasma color scheme with low and high % of FRET positive cells colored purple and yellow, respectively (right).

**Supplementary Figure 14.**
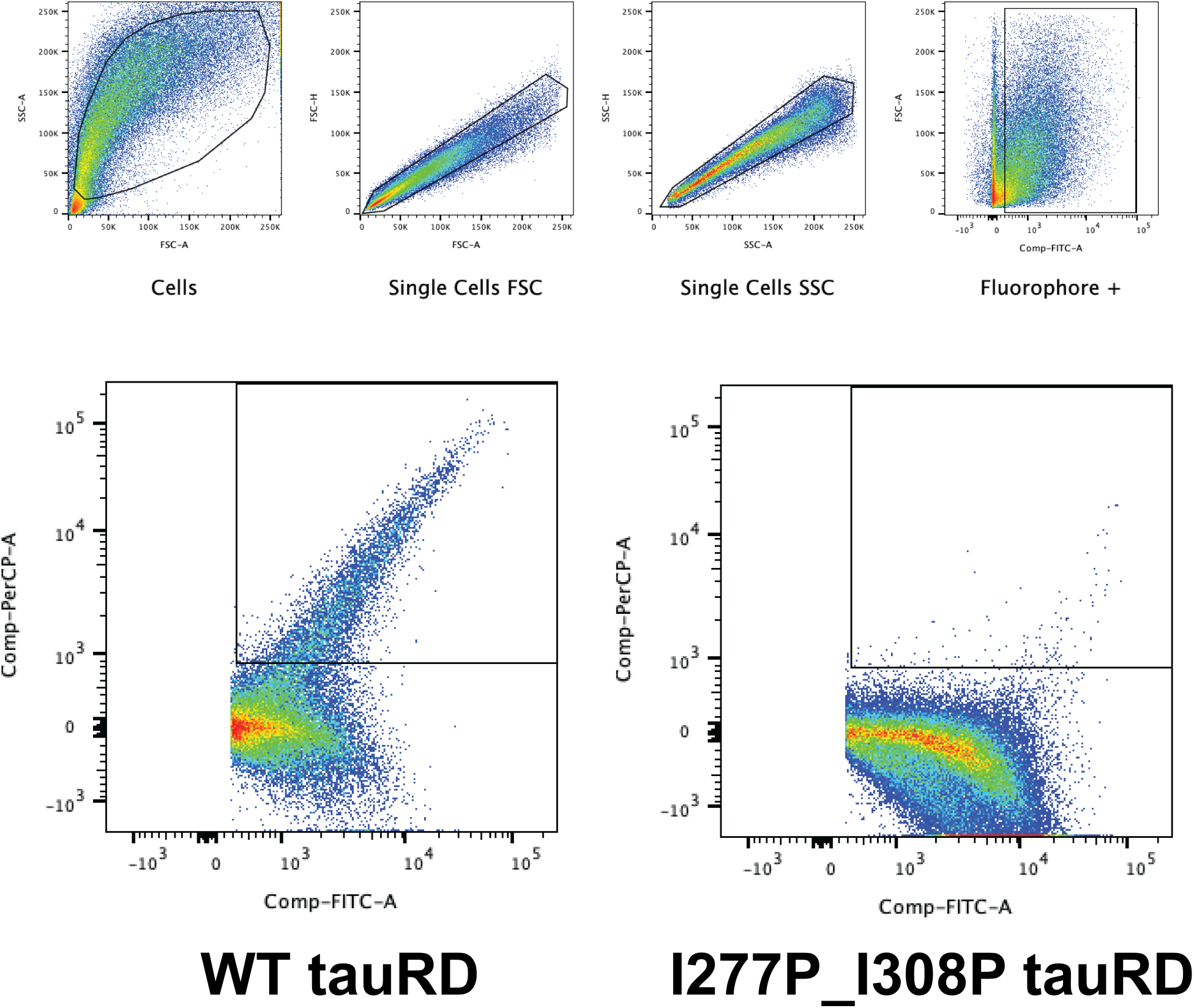
Gating Strategy for Alanine Scanning mutagenesis-based seeding assay with AD, CBD and PSP tauopathy tissues. **(a)** Representative flow cytometry gating strategy and distribution plots used to quantify aggregate-positive cells, showing sequential gating, fluorescence intensity thresholds, and population shifts upon seeded aggregation. Example Gating strategies highlighting populations of FRET positive cells for WT tauRD (bottom left) and negative control I277P_I308P tauRD mutant (bottom right).

**Supplementary Figure 15.**
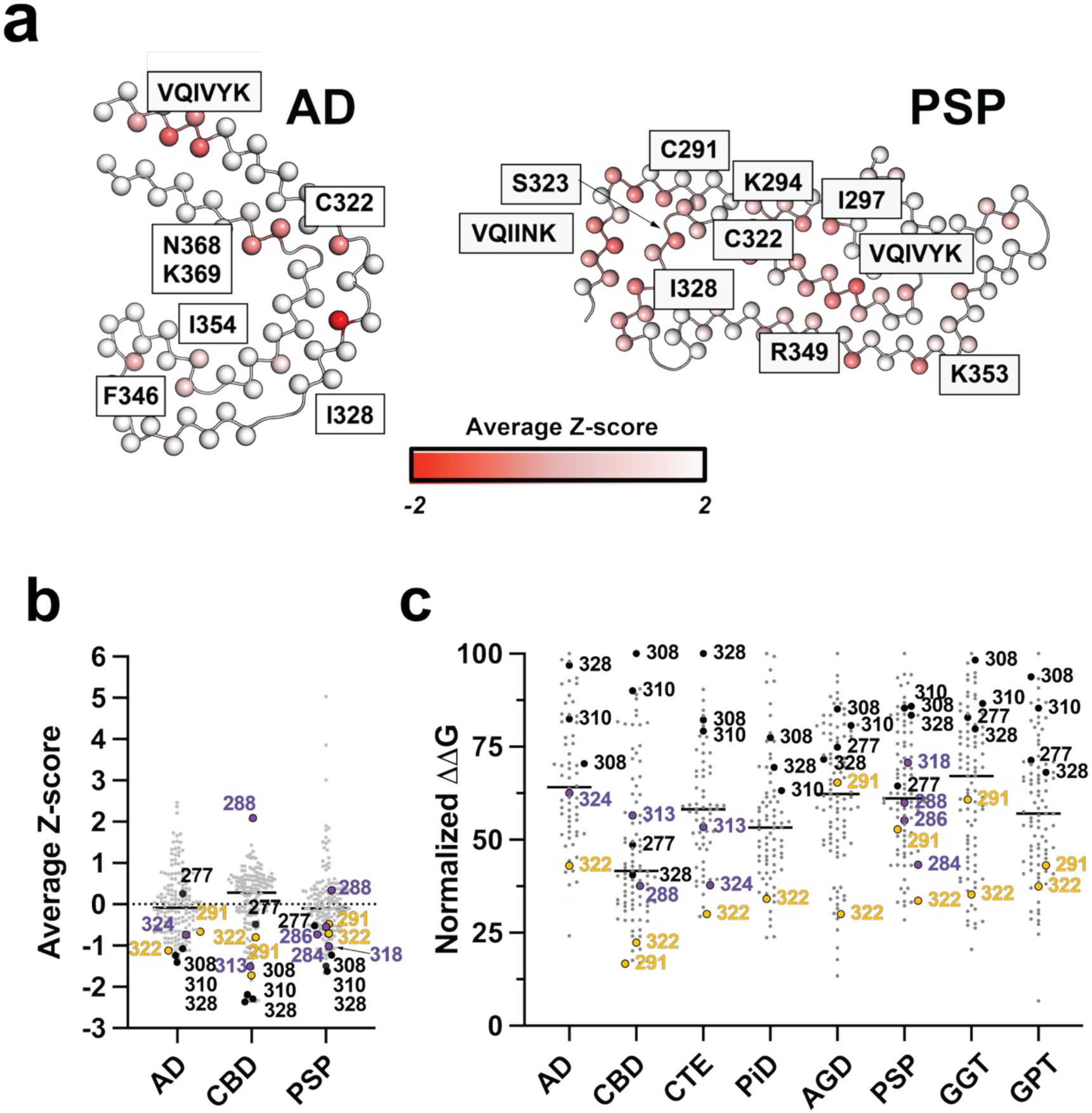
Disease- and seed-specific contributions of tau cysteine residues to aggregation energetics and seeding activity. **(a)** Residue-level maps of average Z-scores mapped onto tau fibril structural models highlighting positions that contribute most strongly to aggregation propensity in Alzheimer’s disease (AD, left) and progressive supranuclear palsy (PSP, right). Core amyloid motifs (VQIVYK and VQIINK) and key residues, including C291 and C322, are annotated. Color scale indicates relative stabilization or destabilization (average Z-score). **(b)** Distribution of average Z-scores for tau repeat-domain residues across disease contexts, emphasizing the prominent contributions of C291 and C322 relative to surrounding positions. **(c)** Normalized aggregation/seeding activity for alanine substitutions across multiple tauopathy-derived seeds, demonstrating disease-dependent sensitivity to mutations at C291 and C322 compared with other residues. Together, these analyses identify conserved cysteines as major determinants of tau aggregation energetics and seeded assembly, with effects comparable to canonical amyloid-forming motifs.

**Supplementary Figure 16.**
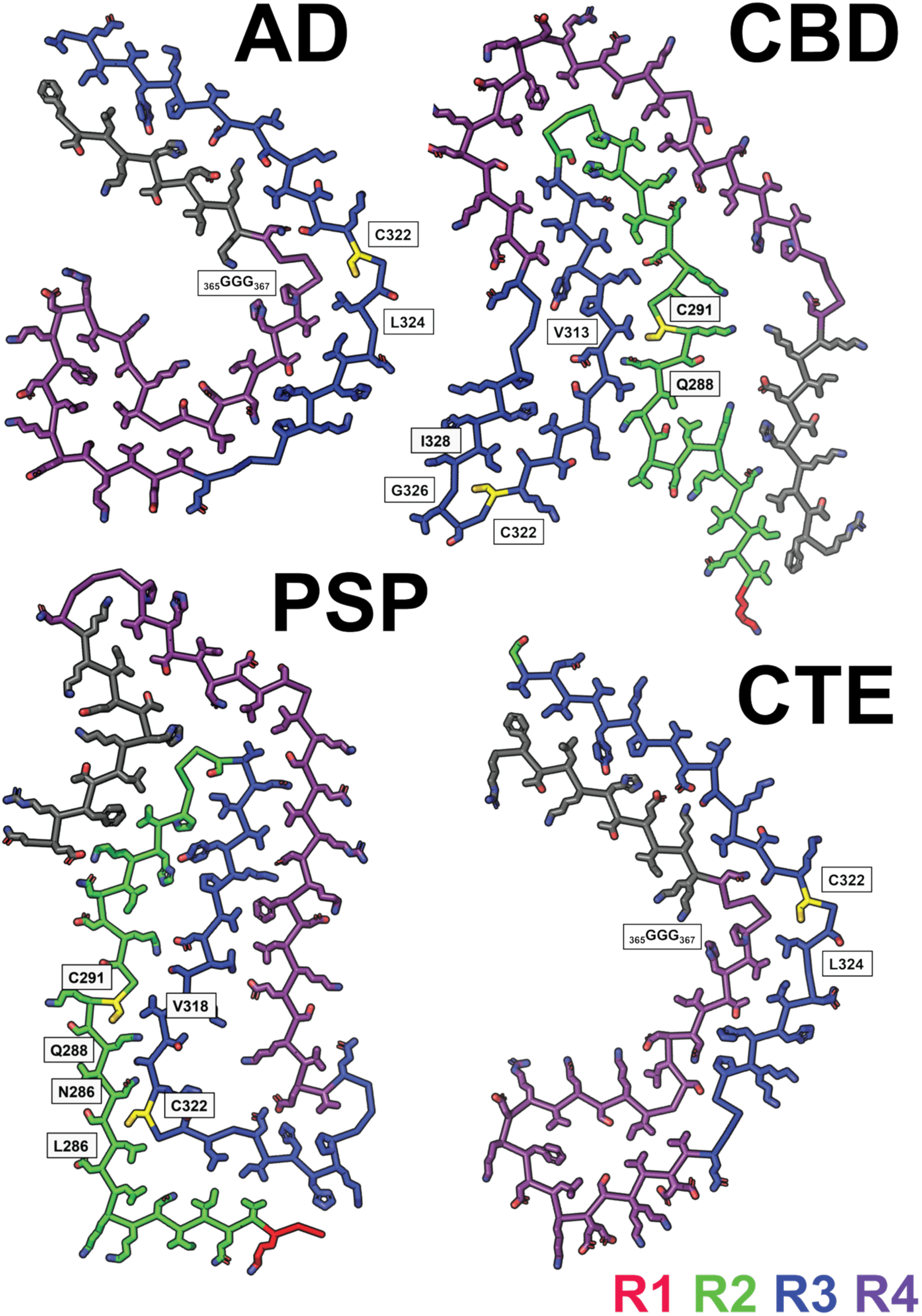
Disease-specific tau fibril architectures highlight distinct positioning of cysteine residues and amyloidogenic motifs. Atomic models of tau repeat-domain fibrils derived from brains of individuals with Alzheimer’s disease (AD) (top left), corticobasal degeneration (CBD) (top right), progressive supranuclear palsy (PSP) (bottom left), and chronic traumatic encephalopathy (CTE) (bottom left). Individual tau repeats are color coded (R1, red; R2, green; R3, blue; R4, purple), with key residues and motifs annotated. Residues are shown as sticks and colored by repeat domain. Cysteine residues C291 and C322 are highlighted in yellow, illustrating their burial or local interactions in each fibril structure.

